# Particular sequence characteristics induce bias in the detection of polymorphic transposable element insertions

**DOI:** 10.1101/2024.09.25.614865

**Authors:** Marie Verneret, Van Anthony Le, Thomas Faraut, Jocelyn Turpin, Emmanuelle Lerat

## Abstract

Transposable elements (TEs) have an important role in genome evolution but are challenging for bioinformatics detection due to their repetitive nature and ability to move and replicate within genomes. New sequencing technologies now enable the characterization of nucleotide and structural variations within species. Among them, TE polymorphism is critical to identify as it may influence species adaptation or trigger diseases. Despite the development of numerous bioinformatic programs, identifying the most effective tool is challenging due to non-overlapping results and varying efficiency across studies. Benchmarking efforts have highlighted some of the limitations of these tools, often evaluated on either real or simulated data. However, real data may be incomplete or contain unannotated TEs, while simulated data may not accurately reflect real genomes. This study introduces a simulation method generating data based on real genomes to control all genomic parameters. Evaluating several TE polymorphic detection tools using data from *Drosophila melanogaster* and *Arabidopsis thaliana*, our study investigates factors like copy size, sequence divergence, and GC content that influence detection efficiency. Our results indicate that only a few programs perform satisfactorily and that all are sensitive to TE and genomic characteristics that may differ according to the species considered. Using *Bos taurus* population data as a case study to identify polymorphic LTR-retrotransposon insertions, we found low-frequency insertions particularly challenging to detect due to a high number of false positives. Increased sequencing coverage improved sensitivity but reduced precision. Our work underscores the importance of selecting appropriate tools and thresholds according to the specific research questions.

## Introduction

Recognized as being among the most important players in the evolution of genomes, transposable elements (TEs) represent a real challenge for bioinformatics approaches to detect them. TEs are repeated sequences present in almost all eukaryotic genomes. They have the ability to move and replicate, forming different families of similar but not always identical sequences. Several types have been described, depending on their structure and their mode of transposition, varying both in genomic distribution and in sequence length (Wicker et al. 2007). For example, LTR-retrotransposons represent sequences of approximately 10 kb but DNA transposons such as MITEs (Miniature Inverted Repeats Transposable Elements) span only a few hundred base pairs. Moreover, TEs are not randomly distributed in the genome since their insertion patterns reflect a balance between selection pressure against their deleterious effects and genetic drift (Bourque et al. 2018). As a consequence, TEs are likely to be found inserted into each other constituting nested insertions, which are particularly difficult to automatically identify (Bergman and Quesneville 2007). In addition, their proportion in genomes can vary greatly, ranging from a few percent as for example in the honeybee (Weinstock et al. 2006) to the major part of the genome as in maize (Schnable et al. 2009). Over the past twenty years, different bioinformatic tools have been developed allowing their annotation in assembled genomes (Lerat 2019). However, the rapid development of new sequencing technologies has made it possible to access numerous data from different individuals or populations in order to characterize the nucleotide and structural variations within a given species. Indeed, a reference genome for a given species is not sufficient to reflect the overall diversity of individuals. In particular, although TEs are generally regulated in a genome to prevent their activity, certain TE families can nevertheless continue to transpose throughout the life of an individual or may be reactivated due to some stress (Di Stefano 2022). It has been proposed that in Drosophila, the transposition rate is comparable to that of the nucleotide mutation rate (Adrion et al. 2017). More recently, according to the TE family, the transposition rate has been shown to be higher with an average of 4.93 × 10^-9^ insertions per site per generation corresponding to a new insertion in each new embryo (Wang et al. 2023). In humans, the most active TEs have a transposition rate of one insertion every 20 births (Cordaux and Batzer 2009). We can thus expect to find variations in the TE insertion pattern between individuals, which constitutes the TE polymorphism. Polymorphic TEs are particularly important to identify since they represent insertions that may be at the basis of species/population adaptation or triggering diseases. For example, numerous polymorphic TEs have been detected in sub-populations of the Chinese white poplar (*Populus tomentosa)* some of them being under positive selection while inserted in genes involved in stress, defense and immune responses (Zhao et al. 2022). In humans, a specific polymorphic TE insertion is associated with the development of the Fukuyama type congenital muscular distrophy (Kobayashi et al. 1998).

In order to search for polymorphic insertions, bioinformatics tools have been developed to answer specific questions and on particular organisms such as Drosophila, human or some plants (Lerat 2019). All these methods follow similar principles in their functioning which consist first in mapping sequenced reads to a reference genome and a set of reference TE sequences. Then two approaches, that can be combined, have been proposed to detect the presence/absence of TEs. The first is to consider discordant read pairs with one read mapping uniquely on a genomic location and the other mapping on different sequences of the same TE family. The second approach considers split reads, *i.e.*, reads overlapping a junction between the genome and a TE insertion, with a part of the read mapping uniquely on the genome while the other part maps on several TE sequences. More than twenty programs have been developed during the past ten years (for an exhaustive list, see https://tehub.org/), which makes it difficult for users to determine which program is the most appropriate or the most efficient. In particular, the results of these programs are often not entirely overlapping (Ewing 2015; Lerat et al. 2019). This makes it more difficult to identify true positives, especially in the case of non-reference insertions, which correspond to insertions not present in the reference genome but present in the analyzed read samples. Several attempts have been previously made to benchmark all these programs (Nelson et al. 2017; Rishishwar et al. 2017; Vendrell-Mir et al. 2019; Chen et al. 2023). These works showed that all these programs often are not as efficient as indicated in their original publication.

However, these evaluations were made either on partial real data or on simulated data without controlling all parameters, or were targeting only particular TE types like for example the approach by Vendrell-Mir (2019). A problem with real data is that they may be only partial or may contain unannotated TE insertions that can blur the results. However, using partially simulated data is also problematic since it usually does not reflect in a realistic manner a real genome and does not allow to control all parameters. For example, the approach used by Rishiwar et al. (2017) consisted in the random insertions of consensus sequences from three human TE families into human reference chromosomes. In the work by Nelson et al. (2017) and Chen et al. (2023), they inserted one single TE from one of the four active families of yeast at positions that are supposed to be biologically sound. These approaches are thus very biased toward the particularities of a single species. Hence, there are still several unanswered questions regarding the underperformance of certain tools, particularly in relation to specific characteristics of the studied genome and the TE sequences themselves that cannot be achieved using real data or simulated approaches used until now.

In this study, we have developed a simulation approach to produce data based on real genomes to allow the complete control of all genomic parameters. Using data generated for *Drosophila melanogaster* and *Arabidopsis thaliana*, we evaluated several TE polymorphic detection tools and investigated different characteristics like the copy size, the sequence divergence, the distance between copies, the GC content of the surrounding genomic regions, the Target Site Duplicate (TSD) size or the TE family that could explain why some insertions are better detected than others. Our results show that only very few of the different tested programs give satisfactory results and that all programs are sensitive to TE and genomic sequence characteristics that slightly differ according to the species considered. As an application case, we used *Bos taurus* real population data to identify polymorphic LTR-retrotransposon insertions. Low-frequency insertions appeared to be more challenging to detect due to a high proportion of false positives. Increasing sequencing coverage improved the sensitivity but at the expense of precision. Our study emphasizes the importance of selecting appropriate tools and thresholds depending on the scientific questions asked.

## Material and Methods

### Genomic data used for simulation

The sequence of the *Drosophila melanogaster* 2L chromosome version 6.18 in GenBank format was obtained from the NCBI GenBank database (accession number: NT_033779). The chromosome sequence is 23,513,712 bp long in which 3,519 genes and 919 TEs are annotated. For *Arabidopsis thaliana*, a GenBank file of the chromosome 1 was generated using the TAIR10 version of the gene and transposable element (TE) annotation in gff format available from the Arabidopsis Information Resource website (https://www.arabidopsis.org/). The chromosome sequence is 30,427,671 bp long in which 7,509 genes and 7,135 TEs are annotated. The sequence of the chromosome 25 from *Bos taurus* was obtained from the GenBank database (version ARS-UCD1.3, accession number: GCF_002263795.2). The chromosome, that is 42,350,435 bp long, contains 1,006 genes but no TEs have been previously annotated. We thus determined the position of endogenous retroviruses (ERV) using RepeatMasker version 2.0.3 with cattle ERV consensus sequences from Repbase version 29.03 (https://www.girinst.org/). ERV insertions from four ERV families were used for the simulations: two class I ERV families (ERV1-1_BT and BtERVF2) and two class II ERV families (ERV2-2_BT and ERV2-3_BT).

### Simulation tool *replicaTE*

We have developed a simulation tool based on real data. This tool is implemented as several python3 scripts that need to be run successively, using as a starting point a GenBank file (Figure 1). In summary, we consider three types of sequences (genes, TEs and intergenic regions). The genes are cleaned up from any TE insertions meaning that any TE inserted inside the genes, given the annotation, are removed from the gene sequences. Intergenic regions are simulated to remove any misannotated TEs and based on the real intergenic regions with respect to their GC content and length. The real characteristics of TE insertions in the genome (number of copies, size of copies, %divergence, etc.) are used to simulate new TE sequences. These TE sequences are randomly assigned to the intergenic regions. Finally, all three parts are reassembled to create a complete simulated genome and a deleted simulated genome in which half of the TE insertions are removed. The tool is available as a git repository (https://github.com/e-lerat/replicaTE). For the simulation of the three species, default parameters for each module were used.

**Figure 1:**
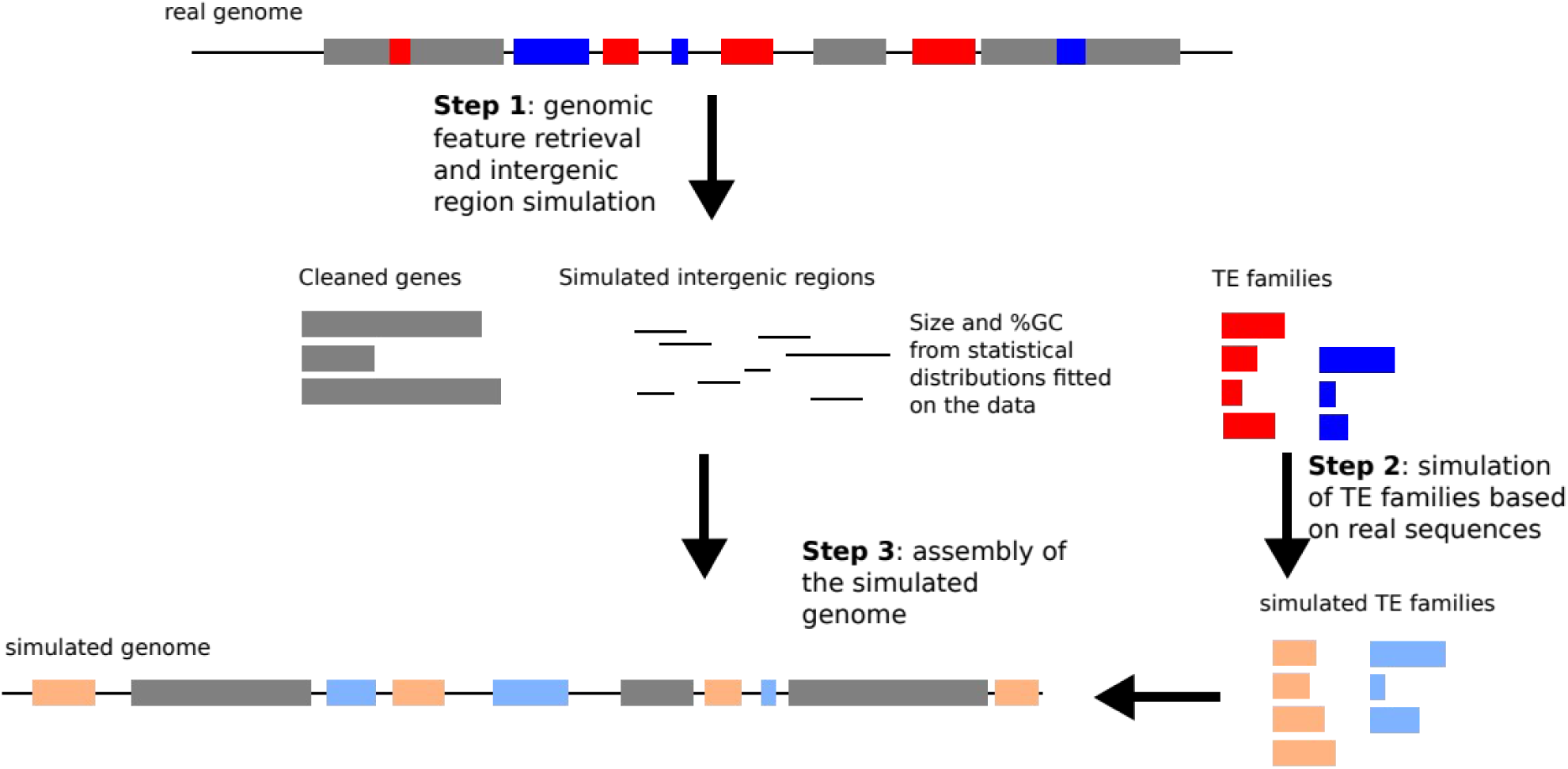
Workflow of the simulation tool *ReplicaTE*. Each step corresponds to the different scripts. In gray are indicated the genes, in blue and red are indicated the TE insertions, a given color corresponding to a given TE family. The simulated TE copies are represented in orange and light blue.

#### deleTE.py

This script allows us to get the characteristics of each element (genes, intergenic regions, and TEs) for the next steps and to generate a simulated genome without TEs. It takes as an input a GenBank file from which it will extract the annotations. It outputs multiple files which can then be used as input by the other codes (see results for the description of these files). The size of the simulated intergenic regions are drawn from an exponentiated Weibull distribution constructed from the computed gene density (number of genes per Mb) with a minimal size of 200 pb. The GC content of the simulated intergenic regions are drawn from a truncated normal law fitted on the observed %GC of the chromosome sequence, with values between 0 and 100%.

#### generaTE.py

This script generates TE copies based on different characteristics (copy number, length, TSD length, strand). It attributes an intergenic region to each copy to be inserted into with the possibility to have nested insertions. For each family, a pool of copy sizes is drawn in a truncated exponential law, with values between 80 bp and 102.5% of the largest sequence of the family to take into account potential small insertions, called the “ancestral” sequence. The sequence divergences of the copies compared to the “ancestral” sequence are drawn from a truncated normal law distribution, with values between 0 and 20% (mean = 10 and standard deviation = 4). By default, the copy number corresponds to the observed copy number in the real chromosome. It is also possible to simulate the copy number. In that case, it is randomly drawn from an exponentiated Weilbull distribution fitted on the data. The TSDs have a length between 0 and 8 bp and are attributed for a given family when the option is specified.

#### inseraTE.py

This script associates the cleaned genes, the simulated TEs and the simulated intergenic regions to produce a genomic sequence. The TE copies are randomly inserted into their attributed intergenic region. The insertion can be ‘normal’ or ‘nested’ (inserted into a previous TE) and multiple nested events can arise. The complete simulated chromosome is provided in fasta format. A “deleted” version is also generated, in which half of the TE copies are not present.

### Short-read simulation

The different tested tools all use short-read sequences as an input. We thus have generated short reads based on either the “complete” or the “deleted” simulated chromosomes using the program ART Version 2.1.8 (Huang et al. 2012). This program produces theoretical reads expected by an NGS technique on a given genome. For this analysis, we generated paired-end Illumina reads of 150 bp (with a fragment size of 300 bp) using three different coverages (10X, 50X and 100X). Only 15X short reads were produced for *B. taurus* to reflect the landscape of the real cattle data coverage in the public databases.

### Polymorphic TE detection tools

Reference and non-reference insertions were detected in the simulated short reads using the either the “complete” or the “deleted” simulated genomes as a reference with the 12 programs included in McClintock2 (Nelson et al. 2017, Chen et al. 2023) in addition to TEPID (Stuart et al. 2016) and Jitterbug (Hénaff et al. 2015) programs. All the programs were run with default parameters. The read alignments on the reference genomes were made using either bwa (Li 2013) and bowtie2 (Langmead and Salzberg 2012) as implemented in McClintock2 with regard to the internal specificity of each tool. TEPID internally uses bowtie2 and yaha (Faust and Hall 2012). In the case of Jitterbug, the read alignments were performed using bowtie2.

### Statistical analyses

All statistical tests were performed using the R software version 3.6.3 (2020-02-29) (R Core Team 2017). The programs were evaluated according to different metrics described below.

Recall (sensitivity): it corresponds to the proportion of True Positives (TP) among all the TE insertions present in the reference genome. It is computed as: 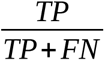

Precision: it corresponds to the proportion of good answers among the predicted TE insertions. It is computed as: 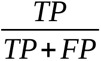

F-score: it corresponds to the harmonic mean of the recall and the precision. It is computed as: 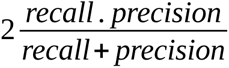

To compute these different metrics, it is necessary to assess the number of TPs among the identified TE insertions proposed by each program, using two homemade perl scripts “test_position_ref.pl” and “test_position_nonref.pl” (available in the git repository). We considered an insertion to be a TP when the program associates the same TE family name and a position that is close to the real position, with a certain margin of error, disregarding the strand of the prediction. More specifically, we considered four different margins of error to determine whether the position was correct or not which are 5 bp, 20 bp, 100 bp and 150 bp. The False Negatives (FN) correspond to insertions present in the reference dataset that were not detected by the program and the False Positives (FP) correspond to predicted insertions that do not correspond to insertions present in the reference dataset.

### False positive rate estimation in real data of *Bos taurus*

Endogenous retroviruses (ERV) insertion detection was performed using TEFLoN (Adrion et al. 2017) with default parameters on 10 WGS short-read data samples from various individuals of *Bos taurus* (accession numbers from SRA database are provided in Supplementary Table S1). A homemade python script (FP_TP_teflon_insertion.py) available in the git repository, was applied to compute the proportion of TPs, FPs and FNs among the identified insertions. Insertions found in common with the reference were considered as TPs or FPs compared to the ERV annotation of the *B. taurus* ARS-UCD1.3 assembly. Insertions found in the samples but not in the reference genome were considered as TPs if they were also present in the variant output file obtain from a variant calling analysis on long-read data from the same samples (accession numbers from SRA database are provided in Supplementary Table S1) using the *call* function of pbsv version 2.6.2 with default parameters (https://github.com/PacificBiosciences/pbsv). In both cases, we considered insertions as TPs if the program also associated the correct ERV family name and with a correct position within 20 bp of error margin.

## Results

### Chromosome simulation and evaluation approach

The simulation tool *replicaTE* was used on the chromosome 2L of *D. melanogaster* and on the chromosome 1 of *A. thaliana* (all generated files are available as supplementary data). The first script, *deleTE.py*, produces different output files. Among them, the “gene_clean_tab.csv” file contains the real genes without any annotated internal TE insertions. The “intergenic_sim_tab.csv” file contains the simulated intergenic regions with their length and %GC. The “stat_TEs_tab.csv” contains a sequence corresponding to the longest real TE sequence (that will be considered as the “ancestral” TE sequence) of a given family that will be used to generate all simulated TE copies and the number of copies for each family, that corresponds to the real number of annotated copies in the considered chromosome. These two last files are used in the second script, *generaTE.py*, to simulate the TE copies. It produces a fasta file containing the simulated sequences (“simulated_TEs.fas”) and a text file (“param_TEs_tab.csv”) containing all the information regarding each TE family (length of each copy, sequence divergence of each copy compared to the “ancestral” TE sequence, the associated intergenic region, the strand and the TSD size). These two files, in addition to the “gene_clean_tab.csv” and the “stat_TEs_tab.csv” files, are used in the third script *inseraTE.py.* It produces, among other files, the two simulated genomes in fasta format and the files “annot_TEs.tsv” and “annot_TEs_del_1” containing all information regarding each TE copy (positions, length, divergence, insertion type (nested or not), strand, TSD size, distances to the closest TE insertions, and the GC content of the flanking genomic regions).

For each chromosome, we thus have all information about the inserted copies in addition to their precise positions. These different parameters will be used to evaluate the tested programs. In particular, we will be able to determine if particular factors relative to the TE sequences (size, distance to other copies, divergence, TSD size) or to their genomic region of insertion (%GC) may have an influence on whether they are correctly detected or not by the tested programs.

In our evaluation approach, the “complete” simulated chromosome and the “deleted” simulated chromosome can be used alternatively as reference genome or as sample genome in order to evaluate the possibility to identify reference / absent insertions or non-reference insertions. Indeed, as described on Figure 2, when using the “complete” simulated genome as a reference, the short reads will be generated using the “deleted” simulated genome, in which half of the TE insertions are missing. This will allow us to evaluate the capacity of the programs to detect both reference and absent insertions. Alternatively, if the “deleted” simulated genome is used as a reference and the “complete” simulated genome is used to generate short reads, then it will allow us to evaluate the capacity of the programs to detect TE insertions not present in the reference.

**Figure 2:**
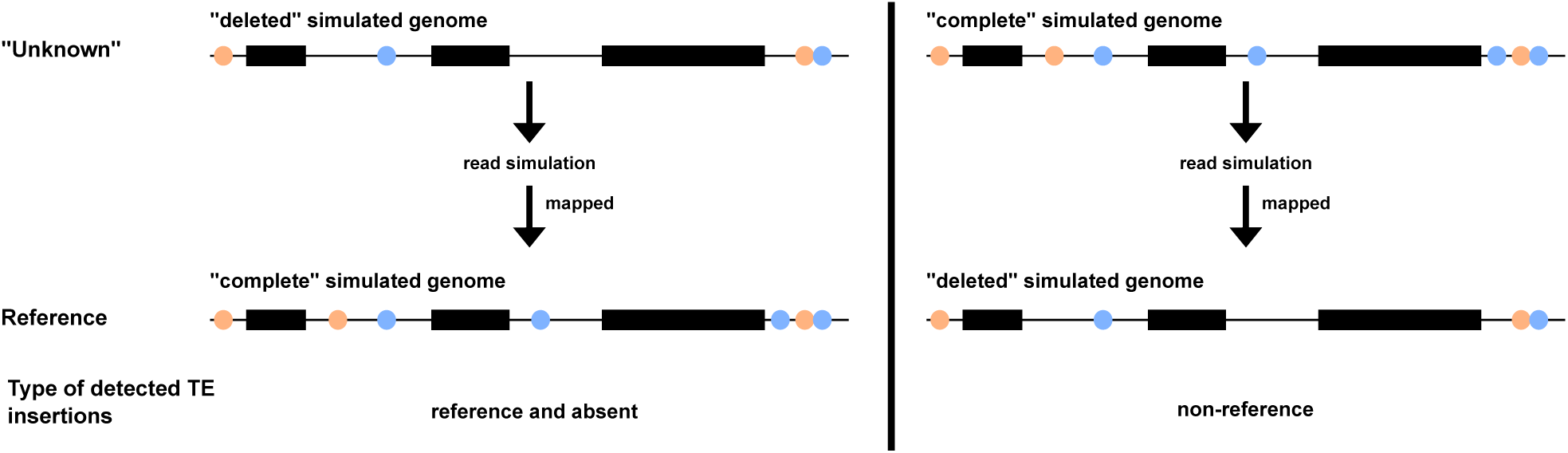
Evaluation approach. The black rectangles correspond to genes. The orange and light blue circles correspond to TE copies from two different families. The reads simulated on the “deleted” simulated genome will be mapped to the “complete” simulated genome, which will allow to identify reference and/or absent TE insertions. The reads simulated on the “complete” simulated genome will be mapped to the “deleted” simulated genome, which will allow us to identify non-reference TE insertions.

Using the *D. melanogaster* and *A. thaliana* data, we have generated two simulated chromosomes for each species. For Drosophila, the “complete” simulated chromosome, which is 23,298,325 bp long, contains 790 TEs whereas the “deleted” simulated chromosome contains 400 TEs. In the case of *A. thaliana*, the “complete” simulated chromosome, which is 37,444,832 bp long, contains 6,324 TEs whereas the “deleted” simulated chromosome contains 3,132 TEs. Knowing exactly the positions and name of each insertion, it is thus possible to compute the number of True Positives (TP), False Positives (FP) and False Negatives (FN) for each program allowing to determine their efficiency. Additionally, since we have all information about the different insertions for which we can control all associated parameters (size, distance, %GC etc.), it will be possible to compare the characteristics of the TP to those of the FN that could indicate any detection bias in the tested programs.

### Tests of the polymorphic TE detection programs

More than 20 programs have been proposed during the last 10 years to identify polymorphic TE insertions. However, many of them were not possible to evaluate in this analysis. Some programs were no longer available to be retrieved. Other programs were not flexible about the reference genome that can be used, unless modifying significantly the source code. We also did not test T-lex3 (Bogaerts-Márquez et al. 2020) since it cannot detect TE insertions present in the sample but not the reference, but only presence/absence of annotated TE insertions in a reference genome.

We have finally tested 14 programs for which it was possible to use customized reference genomes (TEMP (Zhuang et al. 2014), TEMP2 (Yu et al. 2021), ngs_te_mapper (Linheiro and Bergman 2012), ngs_te_mapper2 (Han et al. 2021), PoPoolationTE (Kofler et al. 2012), PoPoolationTE2 (Kofler et al. 2016), RetroSeq (Keane et al. 2013), RelocaTE (Robb et al. 2013), RelocaTE2 (Chen et al. 2017), TEBreak (Schauer et al. 2018), TEFLoN (Adrion et al. 2017), TE-locate (Platzer et al. 2012), TEPID (Stuart et al. 2016), and Jitterbug (Hénaff et al. 2015)). The programs are designed to find non-reference insertions (when compared to a reference genome) and, except Jitterbug, TEBreak and RetroSeq, to find also shared insertions (between a reference genome and a genome under investigation). The programs have been developed on particular organisms but sometimes tested on several of them (human, Drosophila, Arabidopsis, rice, mouse and Daphnia).

#### Detection of reference insertions

We have first evaluated the capacities of the programs to identify reference insertions, that is to say, insertions present in the reference genome and in the genome from which the reads are produced. For the *D. melanogaster* simulated chromosome, it represents 400 insertions and in *A. thaliana*, it represents 3,133 insertions. In Figure 3, the total number of reference insertions found by each program is represented for each species, independently of the identification of true positives (TP). For TEPID, this number has been estimated by subtraction since the program provides information about the absence of reference insertions. As we can see, globally, the increase of coverage has little influence on the total number of reference insertions detected, except for PopoolationTE and PopoolationTE2, which do not find many insertions at 10X. For both species, the TEMP and TEMP2 programs, which have the same results, find far more reference insertions than expected. This may be explained by the way McClintock2 reports reference insertions for these methods since TEMP/TEMP2 find evidence for the absence of reference insertions then McClintock2 computes the complement of the set of “non-absent” reference annotation, which leads to increase the number of reference TE insertions. Other programs find less reference insertions but to a lesser extent in *A. thaliana* (TEPID) and in *D. melanogaster* (TEFLoN for all coverage and PoPoolationTE2 for coverage 50X and 100X). PopoolationTE2, TEFLoN and ngs_te_mapper2 (for *A. thaliana*) and PopoolationTE, ngs_te_mapper2 and TEPID (for *D. melanogaster)* find a number of reference insertions close to what is expected. All the other programs find no or few reference insertions.

**Figure 3:**
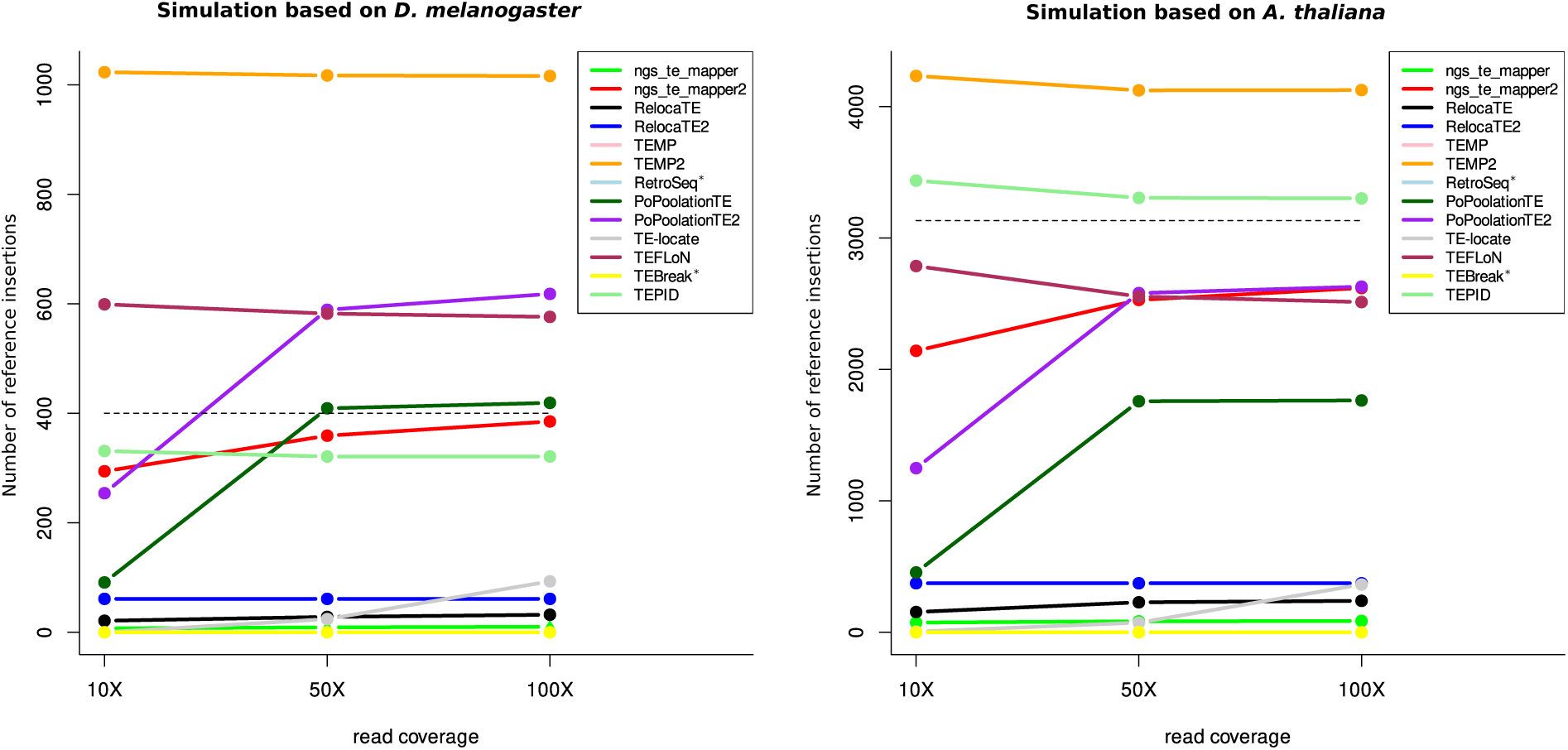
Number of reference insertions detected by each program. On the left panel is represented the number of reference insertions found for *D. melanogaster* and on the right panel for *A. thaliana*, for the three different read coverages. The dashed lines correspond to the expected number of reference insertions in each species; * indicate programs that are not designed to identify reference insertions. Some lines are overlapping and thus are not visible on the figures.

We have then determined among all these insertions the number of False Positives (FP), False Negatives (FN) and True Positives (TP) in order to compute various metrics to evaluate the programs. TPs have been identified according to both the capacity of the program to identify the right TE family and according to the localization prediction with several margin of errors (see Material and Methods). Figure 4 represents the different metrics for both species using a localization prediction with a margin of error of 20 bp (see supplementary figures S1, S2 and S3 for all cutoffs).

**Figure 4:**
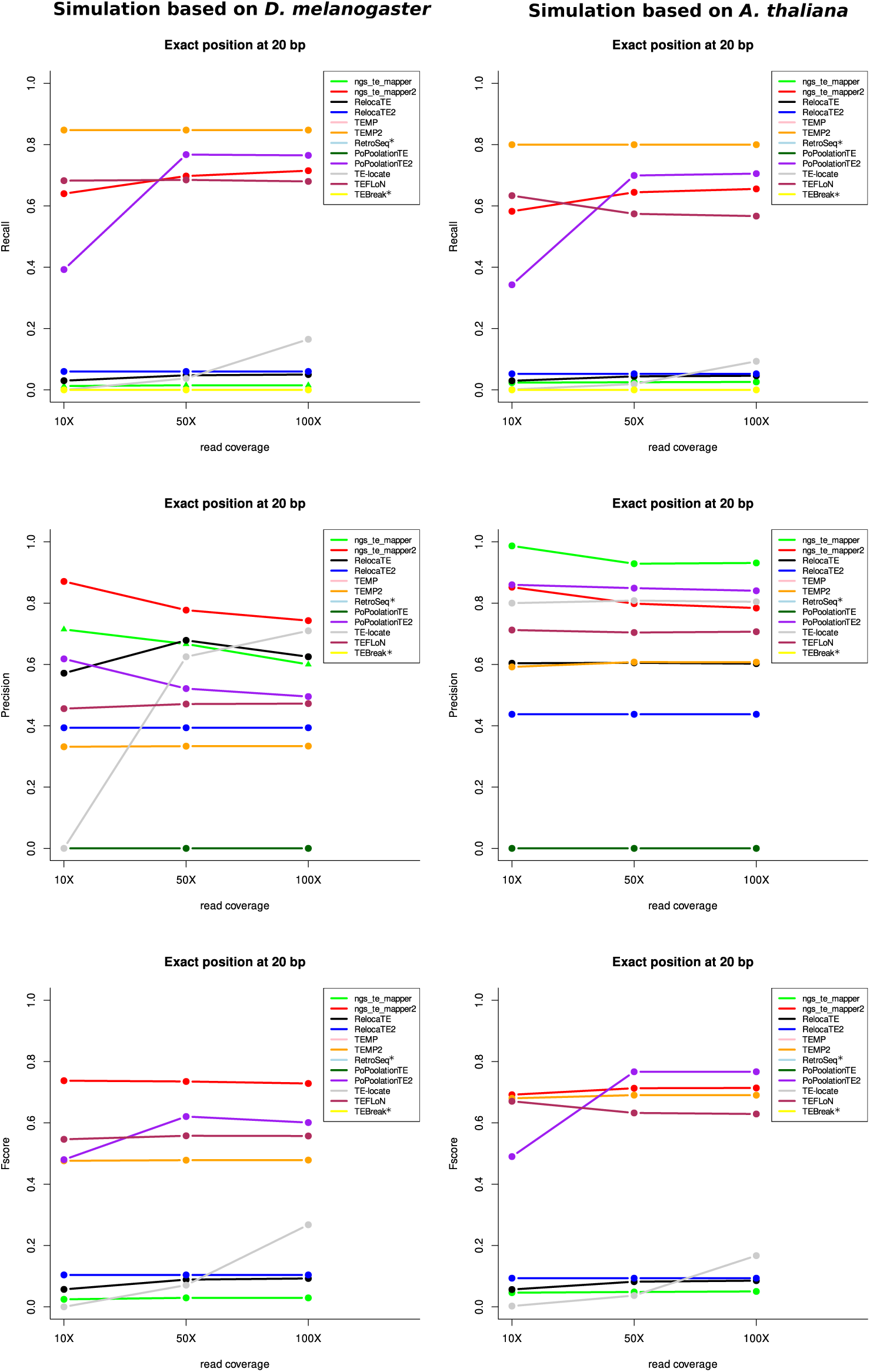
Evaluation metrics for the two species for the three read coverages for the reference insertions with a precision localization of 20 bp; * indicate programs that are not designed to identify reference insertions.

We have retained this particular margin of error since at 5 bp all programs do not perform well whereas at 100 bp and 150 bp the efficiency of the programs is not improved. The *recall* metrics indicate the number of good answers among all the possible predictions. In our case, it indicates for each program the number of TPs among all the reference TE insertions that should be detected. For both species, five programs give the best results for these metrics: TEMP, TEMP2, PoPoolationTE2 (starting at 50X), ngs_te_mapper2 and TEFLoN, with *recall* values of more than 0.5. The other programs find few or no TP among all the TE insertions that can be found given a localization window of 20 bp. The *precision* gives the number of good answers among all the results proposed by the programs. According to the species, the tools do not have the same results. For *D. melanogaster*, ngs_te_mapper2 has the best results for these metrics, whereas it is ngs_te_mapper for *A. thaliana*. In order to take into account both metrics, we have computed the *Fscore*. For both species, five programs give the best results: ngs_te_mapper2, PoPoolationTE2, TEMP, TEMP2, and TEFLoN. However, according to the species, the best program is not the same: ngs_te_mapper2 performs better for *D. melanogaster* when it is PoPoolationTE2 for *A. thaliana*.

We have observed the overlap of TPs between the top four programs for each species (Figure 5). The results show that 66.8% for *D. melanogaster* and 61.9% for *A. thaliana* of the TPs are found by the four programs. Among the remaining TPs, a majority is found in common by at least three programs. Only TEMP/TEMP2 find a significant proportion of unique TPs (3.5% for *D. melanogaster* and 6.1% for *A. thaliana*).

**Figure 5:**
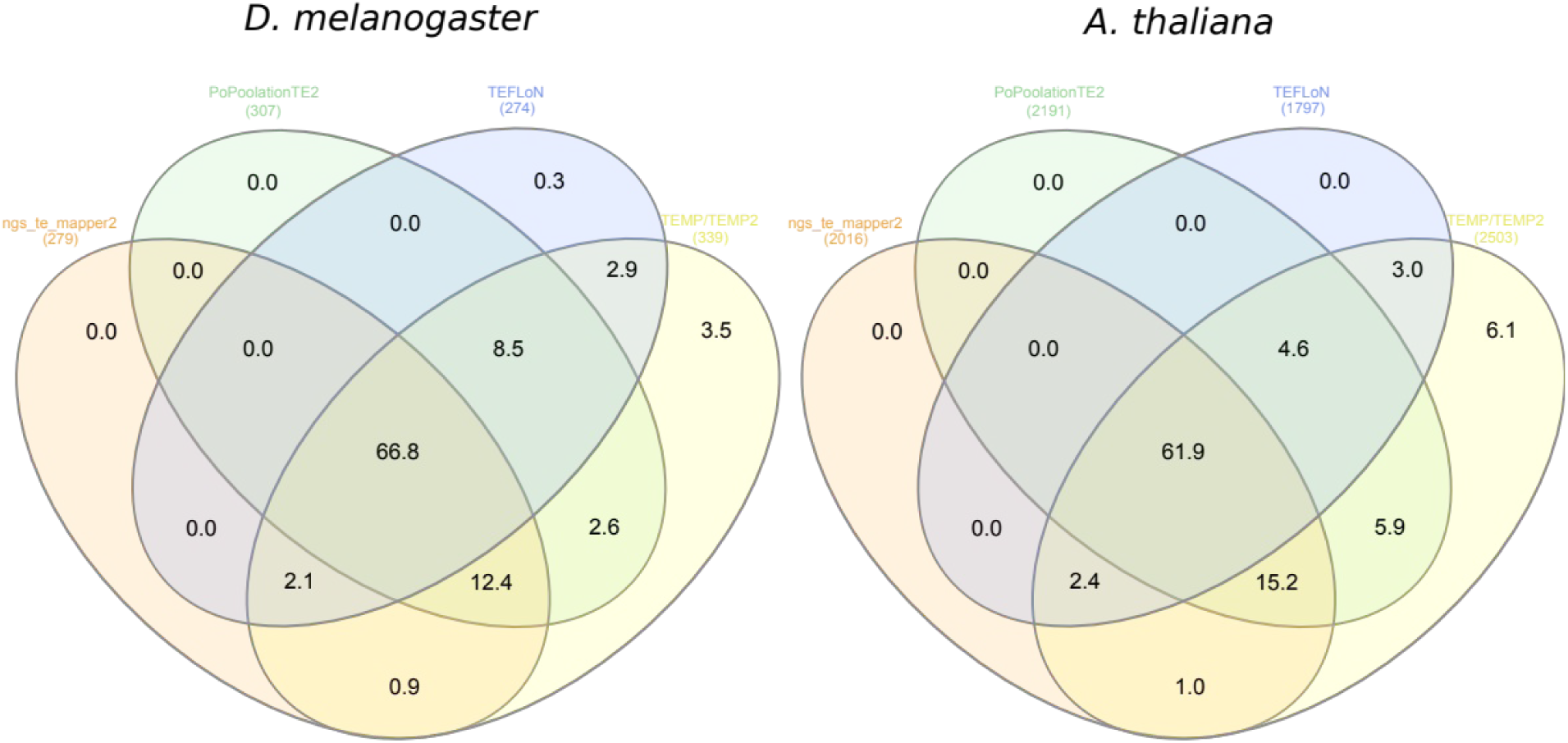
Percentage overlap of TP reference insertions found by the programs having the best *Fscores*.

#### Detection of non-reference insertions

We then evaluated the capacity of the programs to find insertions not present in the reference genome. They correspond to 390 insertions in the simulated *D. melanogaster* chromosome and 3,192 insertions in the simulated *A. thaliana* chromosome. All programs find a total number of non-reference insertions less than what is expected (Figure 6). The sequence coverage has an impact on the total number of non-reference insertions found for the majority of the programs. In particular, a coverage of 10X seems to be insufficient for most programs. Only ngs_te_mapper2 and TEBreak are not very impacted.

**Figure 6:**
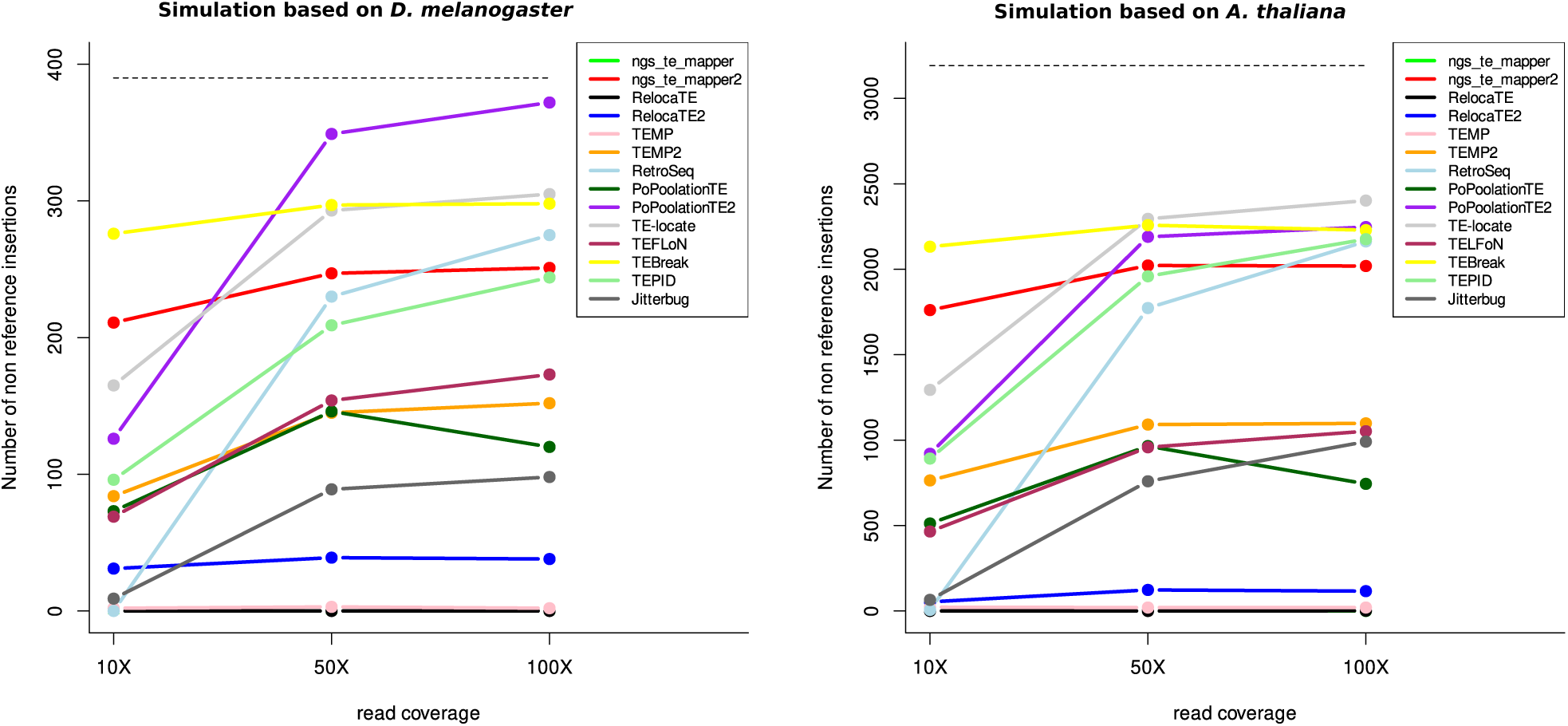
Number of non-reference insertions detected by each program. On the left panel is represented the number of non-reference insertions found for *D. melanogaster* and on the right panel for *A. thaliana*, for the three different read coverages. The dashed lines correspond to the expected number of non-reference insertions in each species.

We have then determined among all the non-reference insertions that are detected which ones are TP according to the same rationale presented above and in the material and methods section. Figure 7 represents the different metrics for both species using a correct localization prediction at 20 bp (see supplementary figures S4, S5, and S6 for all cutoffs). Globally, the recall for each program, and for both species, is not very high, meaning that many TPs are missed by the programs. Three of the programs give the best results considering 50X of coverage (TEBreak, PoPoolationTE2 and ngs_te_mapper2). The *precision* metric on the contrary shows that for most programs, TPs are numerous among all the results produced, especially for *D. melanogaster*. The *Fscore* shows similar results between the two species with four programs having the best results: TEBreak, ngs_te_mapper2, PopoolationTE2 (starting at 50X) and RetroSeq (starting at 50X).

**Figure 7:**
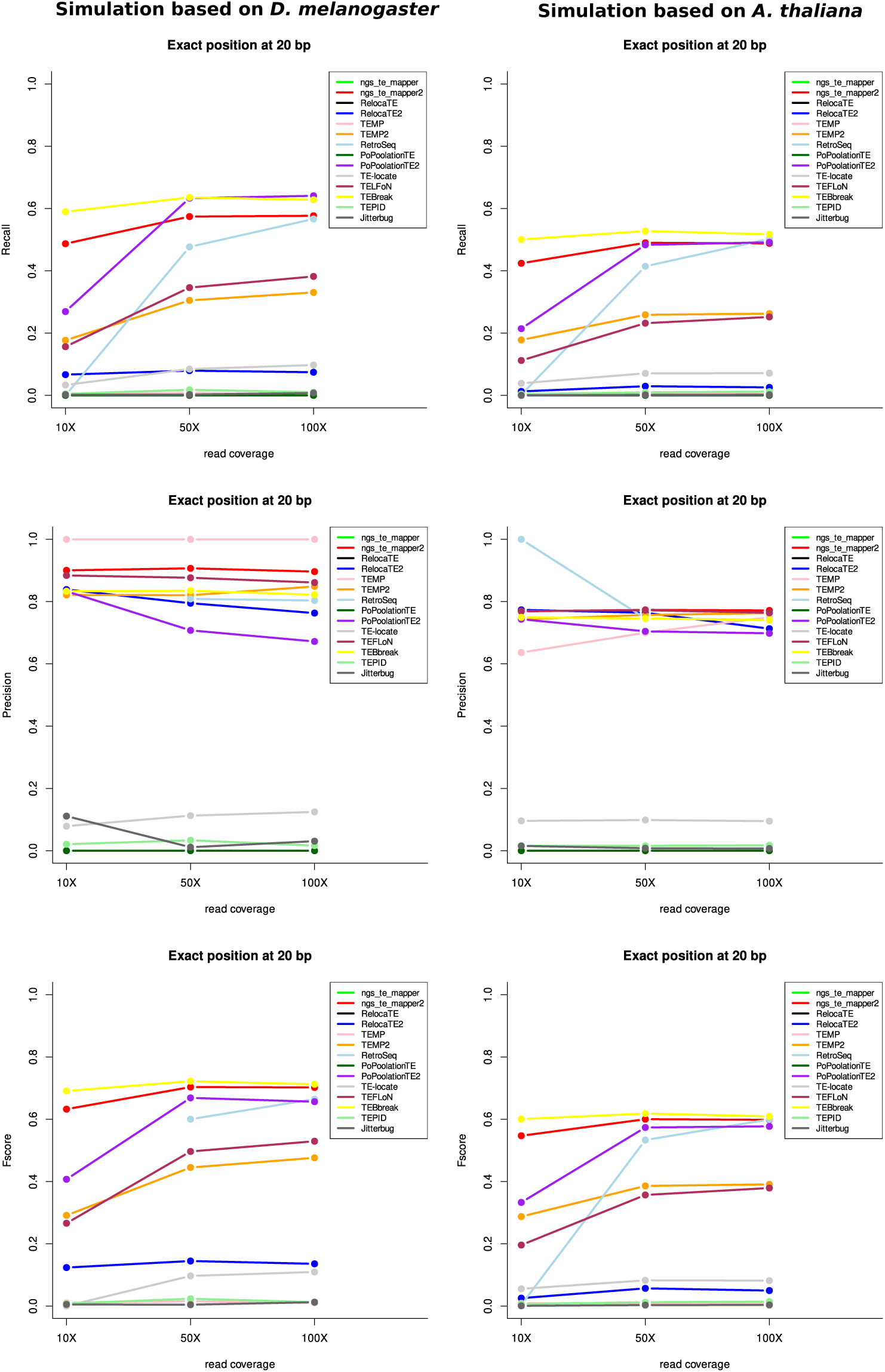
Evaluation metrics for the two species for the three read coverages for the non-reference insertions with a precision localization of 20 bp.

The overlapping of the TP detected by the six best programs accounts for only 10.9% of the TPs for *D. melanogaster* and 10.5% of the TP for *A. thaliana* (Figure 8). PopoolationTE2 and ngs_te_mapper2 each identify 6.5% and 4.4% unique TPs in *D. melanogaster*, and 2.1% and 2.9% respectively in *A. thaliana*. It should be noted that 11.4% of TPs are found by all the programs except TEFLoN in *A. thaliana*. For *D. melanogaster*, 10.6% of TPs are found in common for all programs except TEMP/TEMP2.

**Figure 8:**
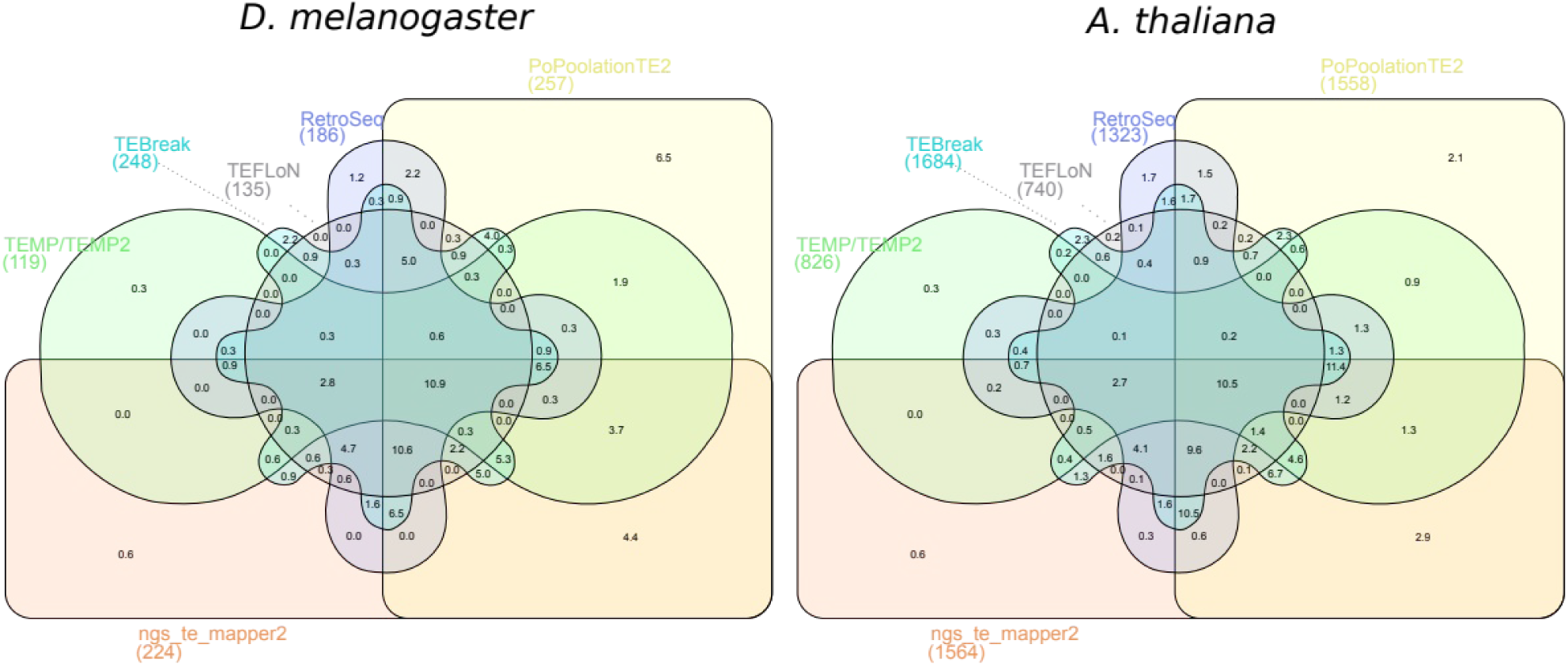
Percentage overlap of TP non-reference insertions found by the programs having the best *Fscores*.

### Comparison of the characteristics between True Positives and False Negatives

Since we know with accuracy the characteristics of all insertions present in the two reconstructed chromosomes for both species, it is possible to determine whether some of them may have an impact on the fact that an insertion is detected or not by the programs. We have considered the results of the programs having the best *Fscores* for a coverage of 50X and with enough identified TPs considering a localization precision of 20bp to allow statistical analyses without bias.

First, we have considered the reference insertions in both species (Table 1 and Table 2, Wilcoxon tests). The results show that the TPs have significantly smaller sizes than FNs for all programs (expected for ngs_te_mapper2 with *D. melanogaster*). Moreover, the distance to the closest TE insertions is also important since it is significantly larger for TPs when compared to FNs, for all programs and for the two species. Additionally, in *A. thaliana*, the %GC of the flanking regions of TPs are significantly more GC rich than those around FNs. To summarize, the programs better detect reference insertions that are small and largely distant from other TE insertions.

**Table 1:**
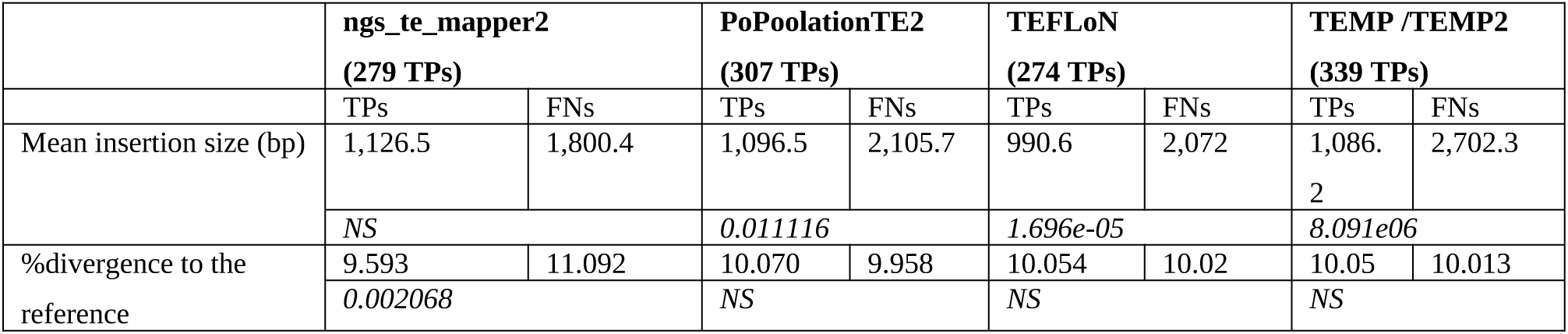
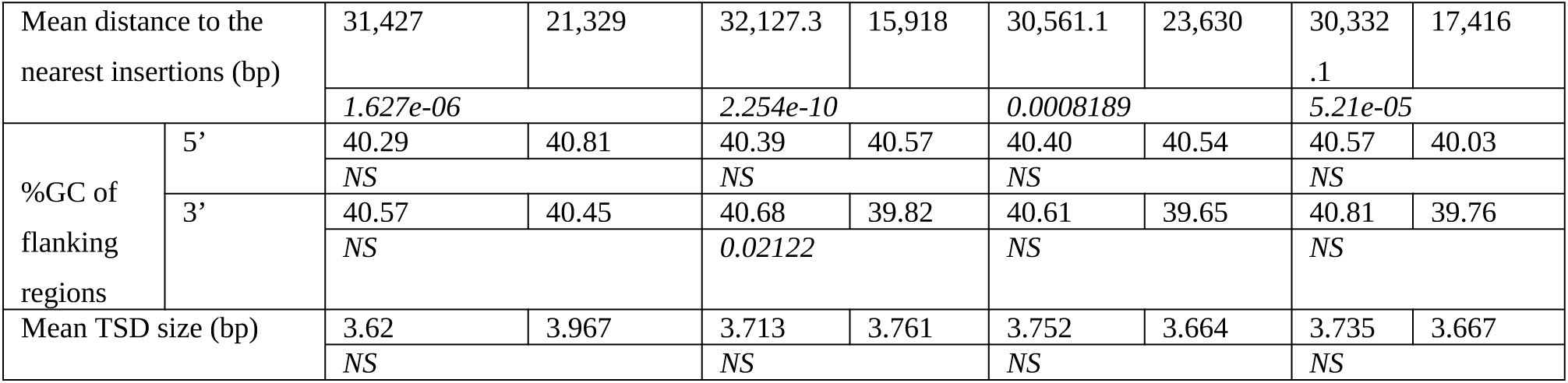
characteristics of TPs vs FNs for reference insertions for *D. melanogaster* (400 reference insertions in total)

**Table 2:** characteristics of TPs vs FNs for reference insertions for *A. thaliana* (3,133 reference insertions in total)

|  |  | ngs_te_mapper2<br>(2,019 TPs) |  | PoPoolationTE2<br>(2,191 TPs) |  | TEFLoN<br>(1,799 TPs) |  | TEMP /TEMP2<br>(2,506 TPs) |  |
| --- | --- | --- | --- | --- | --- | --- | --- | --- | --- |
|  |  | TPs | FNs | TPs | FNs | TPs | FNs | TPs | FNs |
| Mean insertion size (bp) |  | 978 | 1894.7 | 1043 | 1917 | 950.7 | 1781.8 | 1013 | 2465 |
|  |  | <2.2e-16 |  | <2.2e-16 |  | <2.2e-16 |  | <2.2e-16 |  |
| %divergence to the reference |  | 9.70 | 10.33 | 9.91 | 9.93 | 9.81 | 10.1 | 9.88 | 10.1 |
|  |  | 1.435e-05 |  | NS |  | NS |  | NS |  |
| Mean distance to the nearest insertions (bp) |  | 5,684 | 2,455.5 | 5,494 | 2,296.3 | 5,525 | 3,202.5 | 5,156 | 2,059 |
|  |  | <2.2e-16 |  | <2.2e-16 |  | <2.2e-16 |  | <2.2e-16 |  |
| %GC of flanking | 5' | 34.91 | 34.15 | 34.97 | 33.96 | 34.93 | 34.35 | 34.74 | 34.58 |
|  |  | 0.04939 |  | 0.006899 |  | 0.005606 |  | NS |  |
|  | 3' | 35.14 | 33.89 | 35.11 | 33.59 | 35.09 | 34.28 | 34.90 | 33.85 |
|  |  | 5.771e-06 |  | 2.559e-06 |  | 0.002174 |  | 0.03816 |  |
| Mean TSD size (bp) |  | 4.02 | 4.17 | 4.08 | 4.08 | 4.24 | 3.86 | 4.08 | 4.05 |
|  |  | NS |  | NS |  | 0.0003743 |  | NS |  |

In the case of the non-reference insertions (Table 3 and Table 4, Wilcoxon tests), the results show slightly different characteristics. For both species and almost all programs, the percentage of divergence of TPs compared to its ancestral sequence is significantly lower than for the FNs. Again, the distance of TPs to the closest TEs is larger than for the FNs, especially for *A. thaliana* but also for *D. melanogaster* for three programs (TEFLoN, RetroSeq, and TEBreak). Also, the size of TSD is significantly larger for TPs than for FNs for both species and for most of the programs. Finally, in *A. thaliana,* the %GC of the flanking regions of TPs are significantly more GC rich than those around FNs. To summarize, the programs better detect non-reference insertions that are not too divergent from the consensus TE used to identify them (so likely to be recent insertions), largely distant from other TE insertions and with specific TSD size.

**Table 3:**
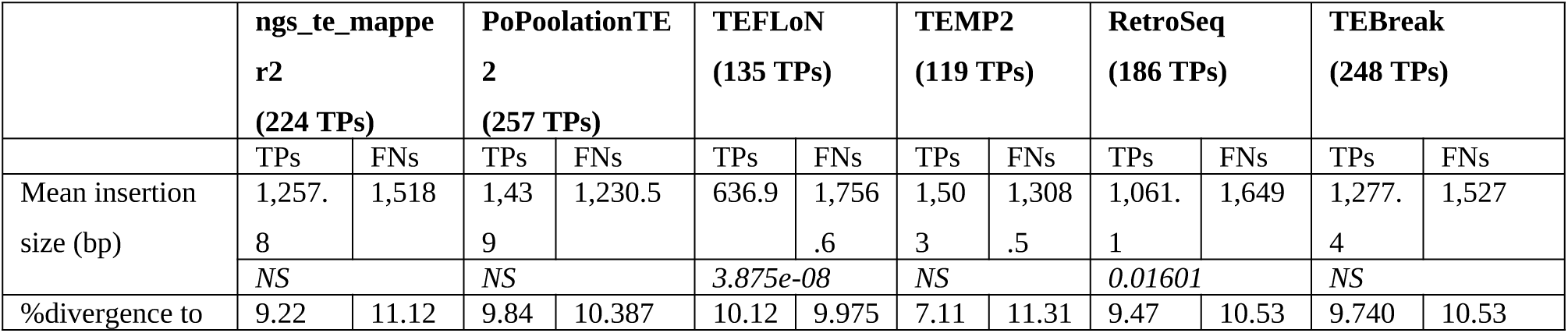
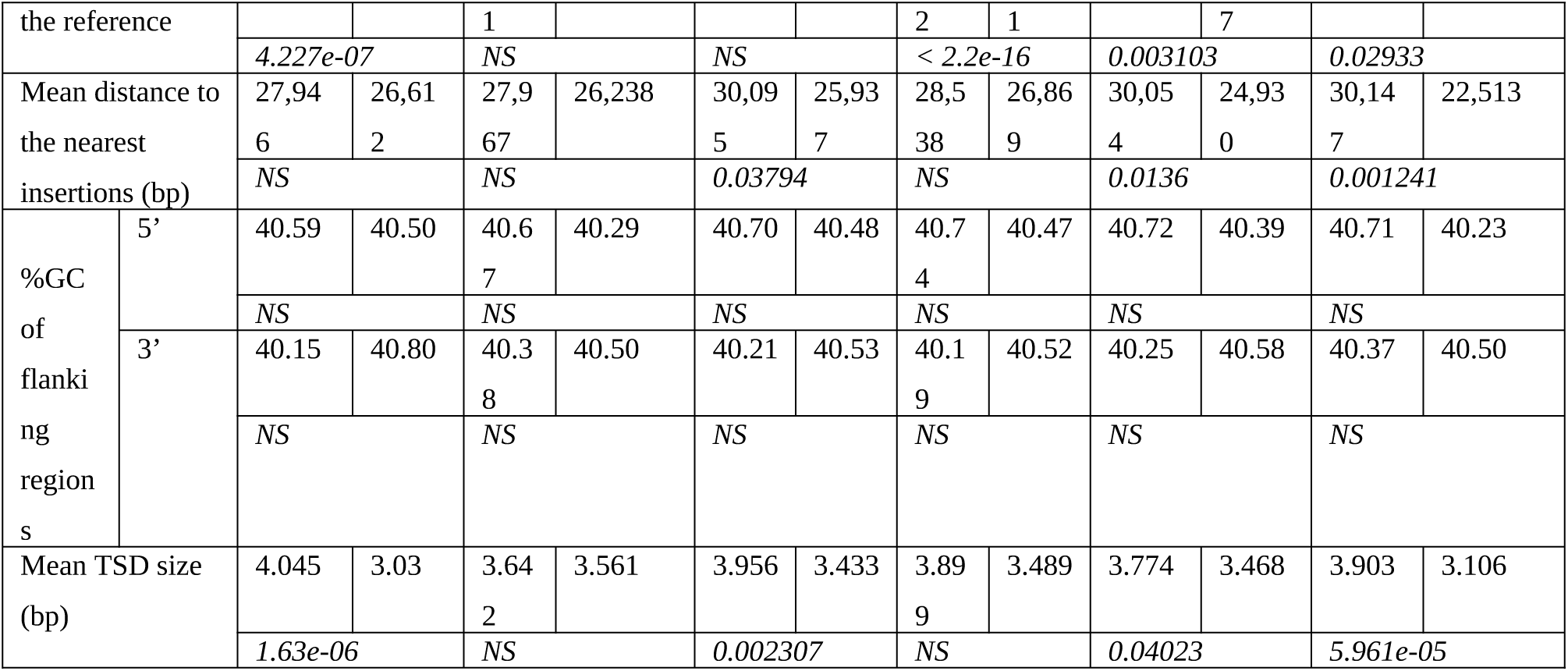
characteristics of TPs vs FNs for non-reference insertions for *D. melanogaster* (390 non-reference insertions in total)

**Table 4:**
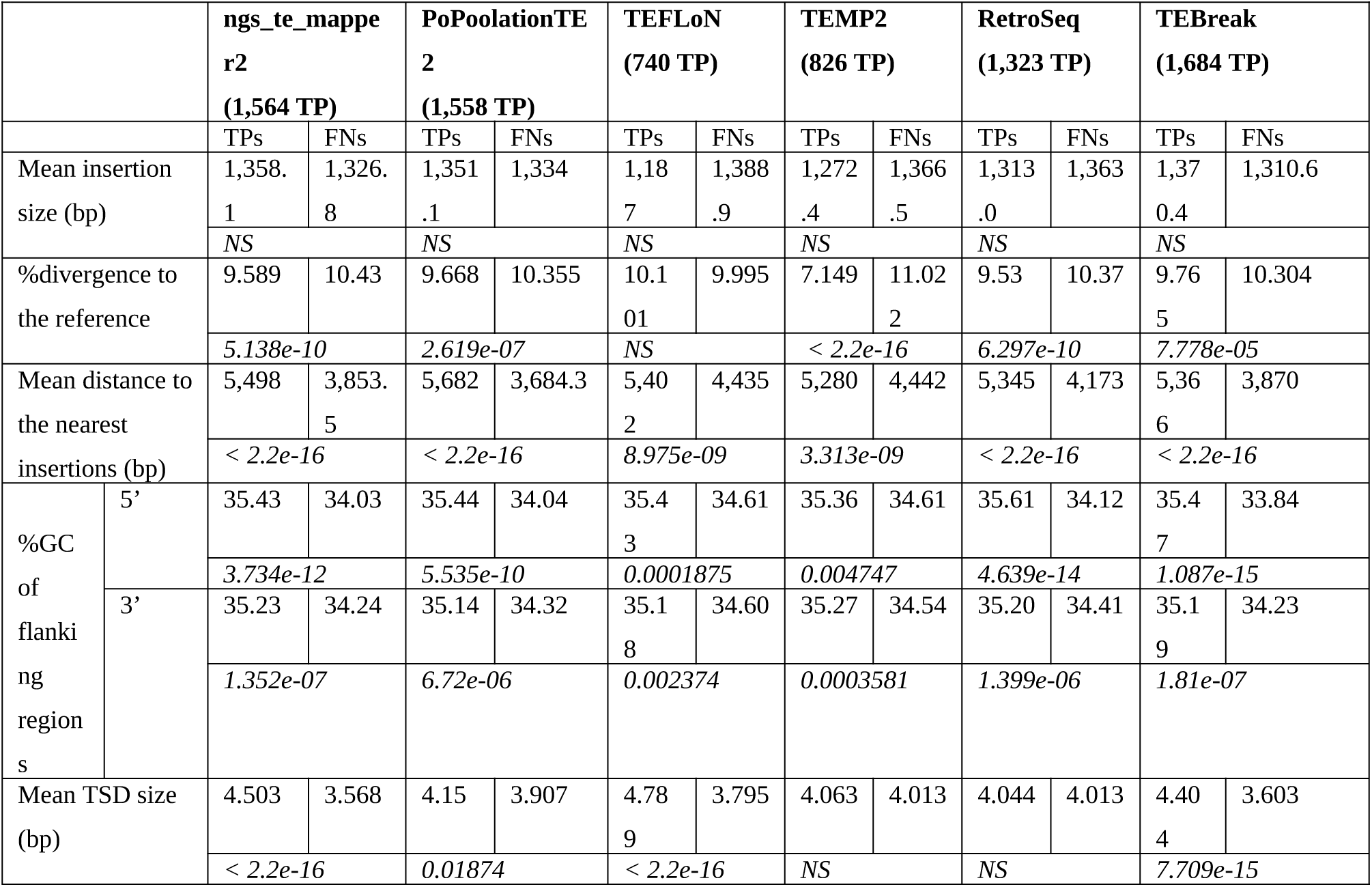
characteristics of TPs vs FNs for non-reference insertions for *A. thaliana* (3,192 non-reference insertions in total)

### Application case: detection of endogenous retroviruses polymorphic insertions in real cattle population data

Although a comprehensive understanding of TEs could have an agricultural interest in improving animal breeding, few TE studies have been conducted on livestock species and more particularly on cattle. We have decided to use cattle as a mammalian genome example to study a subpart of the TEs, the endogenous retroviruses (ERV) insertions. We propose hereafter a workflow to perform such an analysis.

#### Find the best configuration using simulated data

The study of simulated *D. melanogaster* and *A. thaliana* chromosomes has shown that the performance of the programs to detect polymorphic TE insertions are different depending on the studied species. In order to choose the best tool to use, the same pipeline as before has been applied to *Bos taurus* to further detect polymorphic insertions in short-read data. Two simulated chromosomes were generated using *ReplicaTE* from chromosome 25. In this chromosome, 899 random CDS sequences were extracted and 900 intergenic regions were generated. The obtained “complete” simulated chromosome is 25,638,271 bp long including 936 ERVs whereas the “deleted” simulated chromosome contains 474 ERVs. The “deleted” simulated chromosome has been used as a reference and the “complete” simulated chromosome has been used to generate simulated short reads in order to evaluate the capacity of the programs to detect the 462 reference insertions and 474 non-reference insertions. We have determined among the detected insertions the number of False Positives (FPs), False Negatives (FNs) and True positives (TPs) to compute the same metrics as for *D. melanogaster* and *A. thaliana* (see material and methods section).

Figure 9 represents the *Fscore* metric for the detection of ERV reference and non-reference insertions using each of the tested programs (see supplementary figure S7 for *recall* and *precision* metrics). Similar results are found in cattle compared to the other species but the best programs slightly differ. For the reference insertions, TEMP2 and TEFLoN give the best results with a *Fscore* higher than 0.80. For the non-reference insertions, TEFLoN and TEBreak are the two programs giving the best results with respectively a *Fscore* of 0.82 and 0.66. In conclusion TEFLoN appears to be the best performing tool to use on *B. taurus* data.

**Figure 9:**
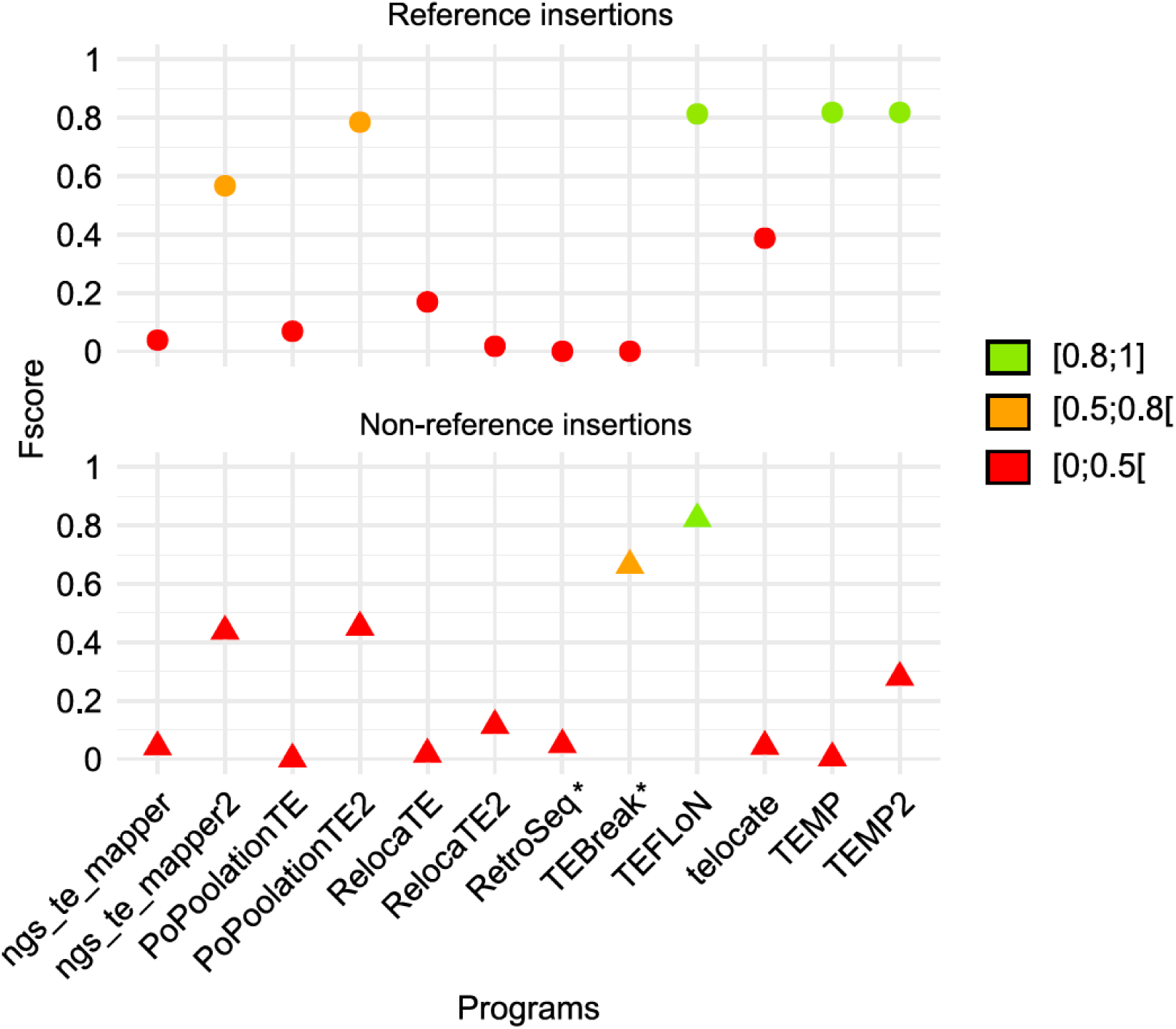
Performance evaluation of McClintock programs on *Bos taurus* simulated data. The detection of the reference and non-reference insertions is represented in the upper and lower panels respectively. INT and LTR consensus were provided separately. The performance of the programs has been evaluated with the *Fscore* metric. * indicate programs that are not designed to identify reference insertions.

Repbase consensus sequences are largely used for TE annotation using RepeatMasker. For LTR-retrotransposons, the LTR sequences and the internal part are often split into two separate sequences. Different types of input sequences have been evaluated to detect ERV insertions in *B. taurus* simulated data using TEFLoN (Figure 10A): i) only the LTR sequences, ii) only the internal sequences, iii) the LTR and internal sequences separately, iv) the LTR and internal sequences concatenated for each ERV family sequence, and v) the LTR, the internal and the concatenated family sequences together to test redundancy. Figure 10B represents the *Fscore* metric for each input sequence (see supplementary figure S8 for *recall* and *precision* metrics). The use of the internal part alone is not working well contrary to other configurations involving both LTR and internal parts. The use of internal and LTR parts separately gives satisfying results for reference insertions but is less efficient for non-reference insertions detection. The input giving the best results is the one with the concatenated family sequences.

**Figure 10:**
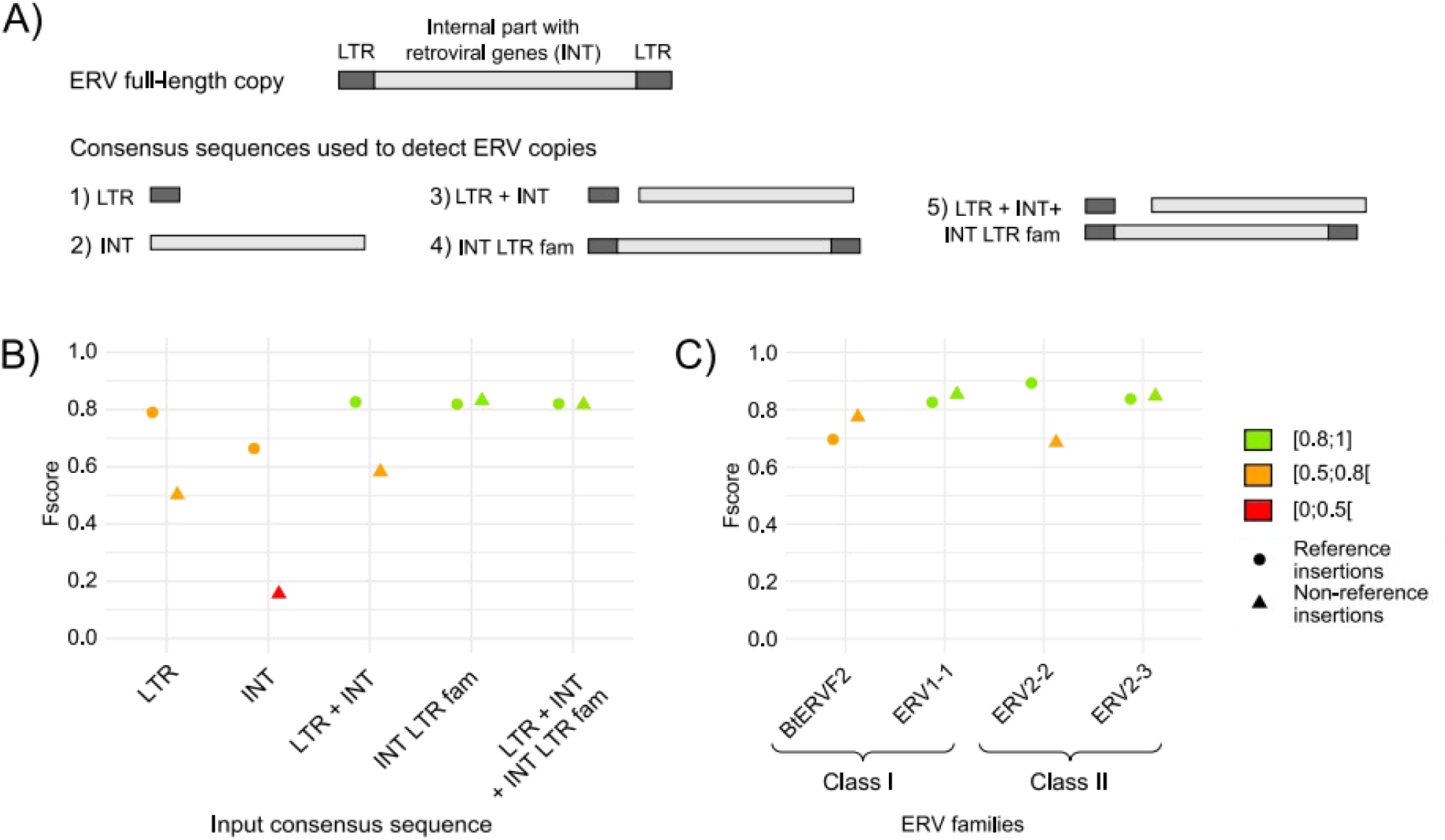
Impact of ERV input consensus sequences and ERV families on insertion detection with TEFLoN on *Bos taurus* simulated data. A) ERV copy genomic structure and the different input consensus sequences tested on cattle simulated data to detect ERV copies, B) *Fscores* (Performance) of TEFLoN with the different input consensus sequences, C) *Fscores* (Performance) of TEFLoN using the detail of each ERV family when using the consensus sequences labeled “INT LTR fam”. The detection of reference insertions is represented with circles and the detection of non-reference insertions with triangles.

We have used four ERV families to generate the simulated chromosomes from ERV class I and II clades. The figure 10C shows how the different ERV families have been detected by TEFLoN. Each family seems to have its own detection characteristics that might correspond to sequence characteristics identified for *D. melanogaster* and *A. thaliana*.

#### Detection of insertion polymorphism in real population data

We have used the previously selected tool TEFLoN to analyze 10 cattle WGS short-read datasets. The detected insertions have been compared to the ERV annotation of the reference assembly and to the output of a variant calling analysis performed on long-read data from the same samples. Figure 11 represents the *Fscores* obtained for these samples and the correlation between the tool performance and the sample short-read depth sequencing. More than 80% of the expected insertions are detected, on average, in the 10 samples. ERV insertions also present in the reference genome are significantly better recognized than the non-reference insertions (Wilcoxon test, p = 1.1e-05). Among the insertions common with the reference, almost no FP are identified. For insertions not present in the reference, almost a hundred of FP are detected representing from 30 to 40% of the non-reference insertions detected in each sample. The tool performances are also more homogeneous between the samples for the detection of reference insertions than for the non-reference ones mainly due the short-read coverage differences across samples. A higher coverage improves the detection of insertions but also increases the detection of FPs (see supplementary figure S9). Furthermore, samples with coverage lower than 10X have a drop in detection rates compared to the others. It appears that 10X is the minimum coverage to reliably detect a sufficient number of ERV insertions. Finally, the comparison between the analysis on simulated and real data shows better results in detecting reference insertions in real data compared to simulated data, with median *Fscores* of 0.97 and 0.81 respectively. On the contrary, TEFLoN is less effective in identifying non-reference insertions in real data compared to simulated data with median *Fscores* of 0.75 and 0.82 respectively (Figure 11).

**Figure 11:**
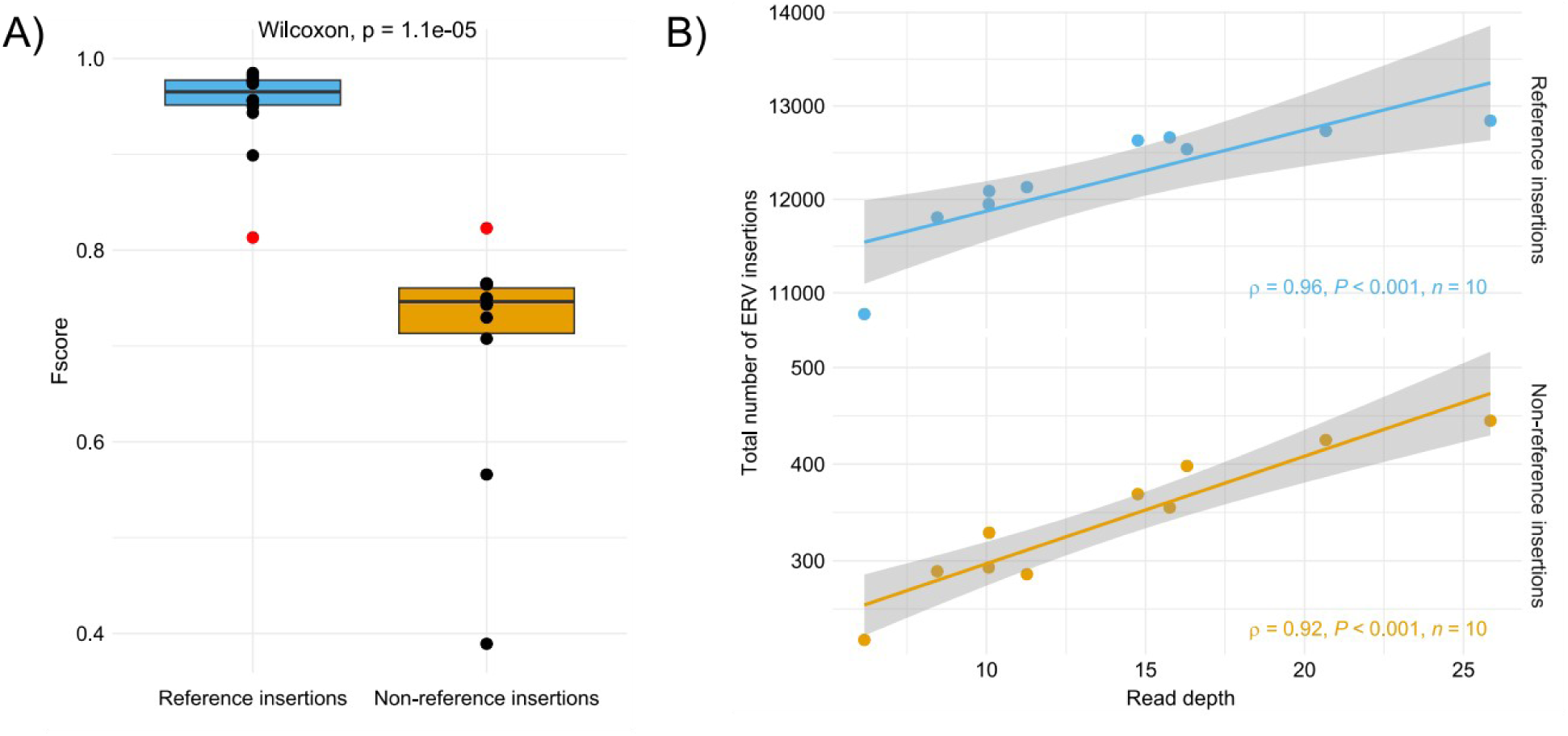
Detection of ERV insertions in 10 *Bos taurus* samples with TEFLoN. A) *Fscores* for the detection of ERV insertions in the 10 samples. The red dots indicate the *Fscores* obtained with TEFLoN on cattle simulated data, B) Impact of the short-read depth sequencing on the number of detected insertions. Depth is computed on trimmed reads mapped on the cattle reference genome.

## Discussion

In this work, we have developed an approach to simulate TE insertions from a known biological context. The data obtained made it possible to test in a reliable and controlled manner 14 programs for the detection of polymorphic TEs. For the first time in the benchmarking of these approaches, it is possible to show why certain insertions are better detected than others by the different programs. Especially, reference and non-reference insertions show different biases. Reference insertions are more correctly detected if they are small and largely distant from other TE insertions. In the case of non-reference insertions, they need to be similar from the consensus or reference TE used to identify them, very distant from other TE insertions and with specific TSD size.

Generally, full data simulation approaches are often used to test polymorphic TE detection programs. They make it possible to perfectly control all the information. The major problem is that often these simulated data do not completely reflect the biological reality. In order to overcome this problem, we have proposed here an approach that uses real data as a starting point and simulates sequences using the biological information of the organism of interest. We thus made the choice to completely simulate the intergenic regions in order to free ourselves from possible bad TE annotations. However, these intergenic regions are not completely randomly generated sequences. In particular, the %GC of these sequences must correspond to what is observed in the analyzed genome. The GC content may be an important factor influencing the detection since it may be a caveat in steps of mapping (Donato et al. 2021). Similarly, the reinserted sequences of the TEs are not the true sequences but come from a real insertion representative of each family contained in the analyzed genome. This allows us to control not only the position of the insertions but also to know with accuracy other information that may play a role on whether an insertion is detected or not. Thus, among the parameters which are controlled, the size of the insertions, the sequence divergence with respect to the reference element, the distance to the closest TEs and the size of TSD are perfectly known for each insertion. It thus allows us to shed light on precise sequence characteristics rather than limiting tests on specific types of TEs, which is sometimes an approach used to benchmark TE polymorphic tools (Nelson et al. 2017; Vendrell-Mir et al. 2019; Chen et al. 2023). Our approach seems to be a good compromise between the use of complete simulation and real but partial biological data with either consensus TE sequences or using only a small set of TE families. However, a number of improvements can be considered with our approach. Currently, only one chromosome is simulated. It could be interesting to simulate several chromosomes and in particular, to generate populations of chromosomes in order to mimic what can be observed in a natural population. Additionally, the tool is currently limited regarding the number of TE insertions that can be inserted. Thus, for a human chromosome for example, the tool works only with a limited number of TE families. The input format of the reference chromosome could be modified to support bed annotation files along with a fasta file containing the chromosome sequence, rather than one genbank file. However, in any case, TE annotations for the reference species are mandatory to allow the different programs to be used to identify reference insertions.

Our results show that all the programs tested here are far from obtaining results as good as announced in their original publication. For some of them, read coverage strongly impacts the ability to find non-reference insertions, as has already been shown (Rishishwar et al. 2017; Chen et al. 2017; Vendrell-Mir et al. 2019) but this is only true up to 50X coverage from which a plateau is reached. Moreover, the results are not as good whether we are interested in the reference insertions (present in the reference genome) or the non-reference insertions (present only in the read samples). Indeed, non-reference insertions are less correctly detected than the reference insertions, an observation that was also made by the only other benchmark evaluating non-reference insertion detection (Vendrell-Mir et al. 2019). Globally, the *Fscores* are better in the first case. However, the values obtained for the best programs do not indicate exceptional performance. Indeed, for reference insertions, the best programs ngs_te_mapper2 and popoolationTE2 have *Fscores* below 0.8. The other programs (PopoolationTE2, TEFLoN, TEMP, and TEMP2) show values around 0.6. For non-reference insertions, the best programs (TEBreak, ngs_te_mapper2, popoolationTE2 and RetroSeq) have values hovering around 0.6. It is important to note that some programs are more successful in finding reference insertions than non-reference insertions, and *vice versa*. TEFLoN, TEMP and TEMP2 show poorer performance in finding non-reference insertions compared to reference insertions. Overall ngs_te_mapper2 and PoPoolationTE2 give consistent results for the two types of insertions. If we compare the results for the two species, there are some notable differences for the detection of reference insertions. Ngs_te_mapper2 gives better results with *D. melanogaster* while the best program is PoPoolationTE2 (at 50X and 100X) for *A. thaliana*. In the case of non-reference insertions, all programs give comparable results for the two species, although working a little less well in the case of *A. thaliana*. It is to note that RelocaTE2 was proposed as the best performing tool to identify non-reference insertions in yeast genomes (Chen et al. 2023), which indicates that the choice of the best performing tool needs to be assessed according to the species under study.

Given that the programs produce many false positives (FPs), an approach allowing to optimize the identification of the true positives (TPs), in the absence of comparison, is to use several tools at the same time to retain only the insertions detected in common. This approach has been used for the analysis of many natural populations of *D. melanogaster* (Lerat et al. 2019). However, the two tools used, popoolationTE2 and TIDAL, showed little overlap in their results. We observed the overlap between the TPs for the best programs identified in this work for reference insertions and non-reference insertions. The proportion of common insertions correctly found by all the programs is quite high in the case of reference insertions since it is almost 70% considering four programs. This proportion is much lower in the case of non-reference insertions with less than 11% for six programs. However, the proportion reach 48.1% for *D. melanogaster* and 58,5% for *A. thaliana* when considering only the results common to TEBreak, ngs_te_mapper2 and PoPoolationTE2, the three programs giving the best results in our benchmark. This remains lower than for the reference insertions. This lack of overlap among the tools has already been observed in other benchmarks (Nelson et al. 2017; Vendrell-Mir et al. 2019). Thus, as proposed by Vendrell-Mir et al. (2019), an approach consisting of using several tools at the same time to optimize the number of TPs must be limited to a few tools at a time. Even with this method, it is important to take into account that some information will be inevitably lost and that the number of polymorphic TE insertions will be underestimated. Another possibility would be to consider evolutionary and biological contexts as it was used before (Manee et al. 2018).

With our approach, it was possible to compare the characteristics of the True Positives (TPs) compared to those of the False Negatives (FNs), *i.e.* the insertions which are missed by the programs, a point that has never been assessed previously by the other benchmark analyses. The goal was to determine if there are biases inherent in the sequences preventing their detection. Between reference and non-reference insertions, some differences appeared. In particular, reference insertions correctly detected tend to be smaller than those not detected by the programs. That would indicate that degraded or small size types of TEs will be better detected as reference insertions. This observation is consistent with the fact that MITE reference elements were better identified than LTR-retrotransposon reference elements (Vendrell-Mir et al. 2019) since MITE elements are shorter than LTR-retrotransposons. On the contrary, the non-reference insertions are better detected when their divergence compared to a reference element is low. Then, recent insertions will be better detected. Although this could be enough to identify recent events, it remains that some of the non-reference insertions may be ancient. These particular insertions would be missed by the different programs. In their original manuscript, almost all programs acknowledge the fact that they cannot detect nested insertions. This is confirmed by our analysis for both types of insertions since TPs present significantly larger distances to the nearest TEs than FNs. Globally, the same bias appears between the two explored species. However, for *A. thaliana*, we observed that the GC content of genomic regions surrounding the insertions also play a role in whether they are detected or not by the program. This species has globally AT rich intergenic regions (DeRose-Wilson and Gaut 2007). We observed that the insertions are better detected when the genomic regions are less AT rich. Since TEs are known to be also AT rich (Lerat et al. 2002; Boissinot 2022), they may be better identified when their base composition is more different from the surrounding genomic regions. We also observed for the detection of non-reference insertions that the size of the TSD is important. Since these sequences may not be well conserved, it may prevent the detection of many insertions.

The case study provided here, focusing on *B. taurus,* allowed us to identify important criteria that should be considered before performing studies on polymorphic TEs in real population data. The choice of the program is crucial and depends on the analyzed species. Indeed, the best identified tool to use on this species is not the same as for *D. melanogaster* and *A. thaliana*. Therefore, it is essential to first perform tests on simulated data built with specific elements from the species of interest to identify the most suitable tool(s) to use. The different programs were all used through the McClintock pipeline (Nelson et al. 2017; Chen et al. 2023) which has a significant advantage to allow the use of multiple tools simultaneously, prevents difficulties in program installation and ensures standardized results. It is also important to carefully select the type of consensus sequences, especially for LTR-retrotransposons. For these elements, usually the LTR and the internal parts are separated in distinct consensus sequences. The re-association of the LTR sequences and the internal parts of a given family is thus necessary and require an in-depth annotation of the reference genome.

Here, we demonstrated the importance of testing a tool also on real data before launching a large-scale population analysis. Even though our study was limited to 10 samples, the genomic characteristics and TE content reflected the reality. The results obtained on real data were different compared to the simulated data, with a better detection of the reference insertions but a less effective identification of the non-reference insertions. This difference is mainly due to the total number of ERV insertions. In the simulated data, half of the total insertions were insertions not present in the reference, whereas they constituted approximately 2% of the insertions in the real data. It appears that detecting non-reference insertions is easier when they represent a larger fraction of the genome of interest.

We showed that non-reference insertions were overall more challenging to detect than the reference ones. Moreover, assessing insertions absent from the reference genome in real samples is challenging because we do not know what to expect, making it difficult to determine whether an insertion is a true or false positive. In our analysis, we used variant calling results obtained from long-reads sequencing data. However, this approach might also miss some insertions, raising questions about its reliability as a reference. Nevertheless, it provides results from two distinct methodologies, ensuring the identification of TPs, even if some are missed.

In conclusion, most of the tested tools do not achieve extraordinary results. There are several biases that prevent them from detecting certain insertions. In addition, the FP rate is particularly high for some tools. Therefore, it is advisable to use a small number of programs simultaneously to optimize the detection of real insertions while keeping a critical perspective on the results.

## Acknowledgments

This work was performed using the computing facilities of the CC LBBE/PRABI and of the IFB-cloud. We thank the SeqOccin project and the Get-Plage platform (https://get.genotoul.fr/) for sharing the bovine data set, and Mekki Boussaha (G2B team, INRAE Jouy-en-Josas) for sharing the associated alignments computed by his team. We thank Carole Lampietro, Claire Kuchly and Caroline Vernette (Get-Plage) for their help with the submission of the sequencing data to SRA. We thank Caroline Leroux (IVPC) and Vincent Navratil (PRABI-Doua) for useful discussions about this work. JT and TF were supported by the INRAE GoatRetrovirome grant for this project. MV PhD fellowship was funded by ANR, grant ANR-22-CE35-0002-01.

## Supplementary data

**Supplementary files:** Output files produced by *ReplicaTE* on the two species *D. melanogaster* and *A. thaliana*.

**Figure S1:** Recall metrics for *D. melanogaster* and *A. thaliana* for the three read coverages for the reference insertions according to the different precision localization tested.

**Figure S2:** Precision metrics for *D. melanogaster* and *A. thaliana* for the three read coverages for the reference insertions according to the different precision localization tested.

**Figure S3:** *Fscore* metrics for *D. melanogaster* and *A. thaliana* for the three read coverages for the reference insertions according to the different precision localization tested.

**Figure S4:** Recall metrics for *D. melanogaster* and *A. thaliana* for the three read coverages for the non-reference insertions according to the different precision localization tested.

**Figure S5:** Precision metrics for *D. melanogaster* and *A. thaliana* for the three read coverages for the non-reference insertions according to the different precision localization tested.

**Figure S6:** *Fscore* metrics for *D. melanogaster* and *A. thaliana* for the three read coverages for the non-reference insertions according to the different precision localization tested.

**Figure S7:** Recall and precision metrics of the different tested programs on *Bos taurus* simulated data.

**Figure S8:** Impact of the structure of the ERV input consensus sequences (panel A) and ERV families (panel B) on recall and precision metrics using TEFLoN on *Bos taurus* simulated data. The detection of reference insertions is represented with circles and the detection of non-reference insertions with triangles.

**Figure S9:** Performance of TEFLoN and impact of read coverage in the detection of ERV insertions in 10 *Bos taurus* samples. A) Recall and precision metrics, B) Number of TP and FP according to the short-read depth sequencing. Depth was computed on trimmed reads mapped on the cattle reference genome.

**Supplementary Table S1**: Accession numbers of 10 WGS short-read data samples of *Bos taurus*.

**Supplementary file 1**: command lines used to run Jitterbug and TEPID.

**Supplementary file 2**: command lines for the pre-processing of long and short read data.

**Supplementary file 3**: TE annotations for *B. taurus*.

