## Supplementary figures and images for "Particular sequence characteristics induce bias in the detection of polymorphic transposable element insertions"

### observed_intergenic_GC_content_distribution.png

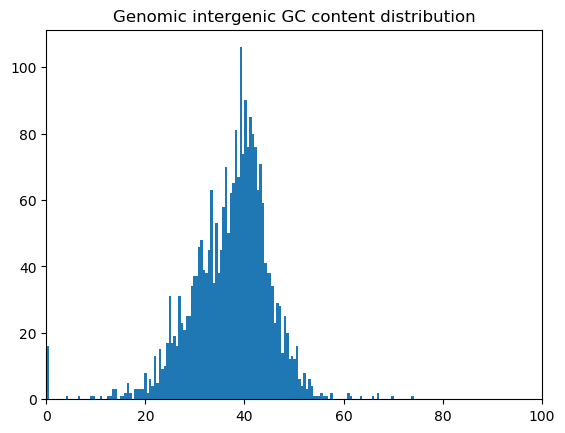

### observed_intergenic_GC_content_distribution.png

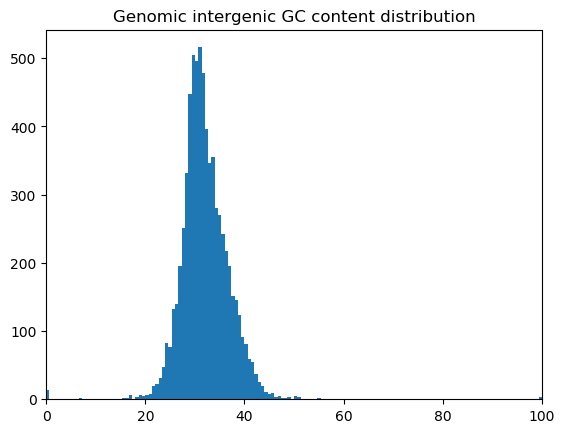

### observed_intergenic_length_distribution.png

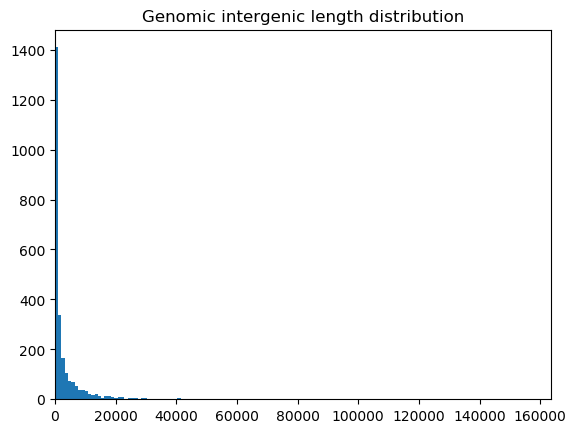

### observed_intergenic_length_distribution.png

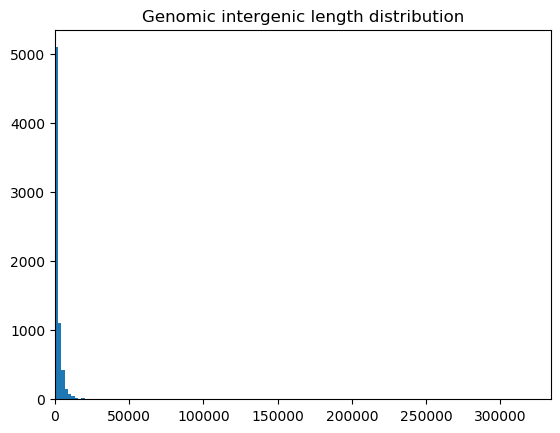

### simulated_intergenic_GC_content_truncated_normal_distribution.png

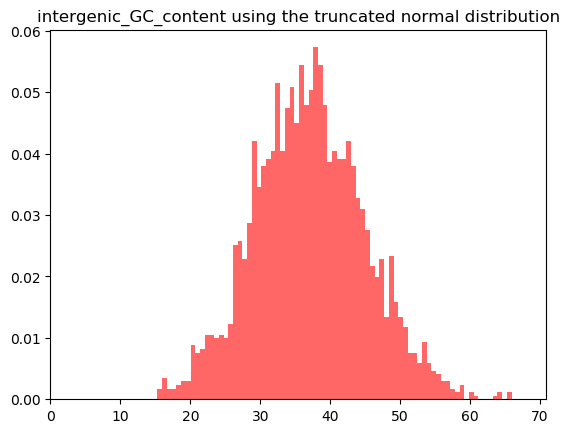

### simulated_intergenic_GC_content_truncated_normal_distribution.png

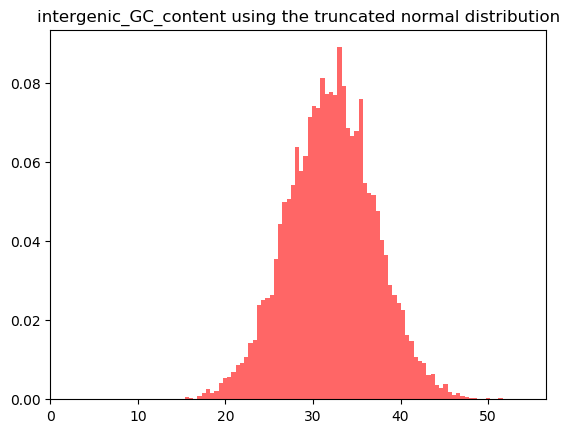

### simulated_intergenic_length_Weibull_distribution.png

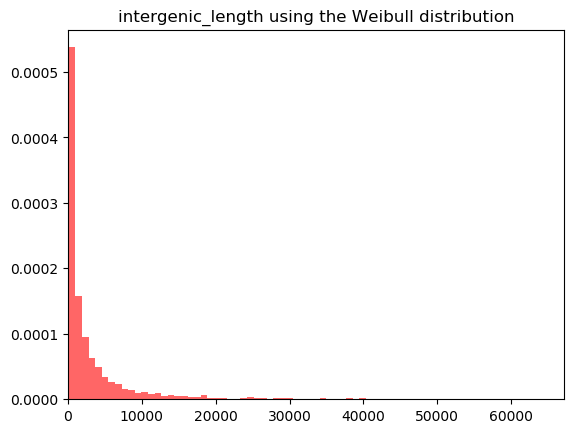

### simulated_intergenic_length_Weibull_distribution.png

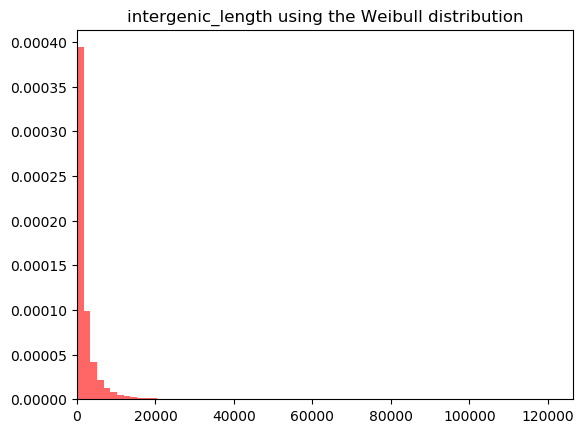

### simulated_TE_copy_divergence_truncated_normal_distribution.png

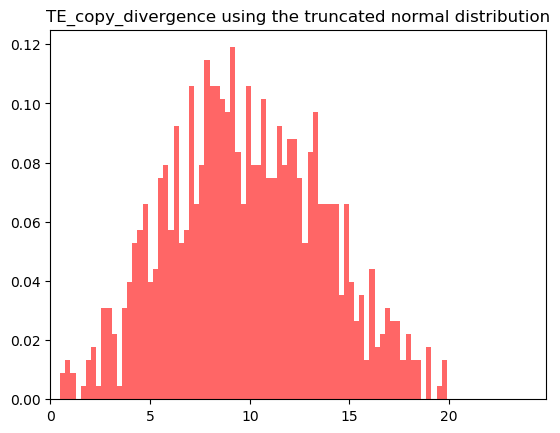

### simulated_TE_copy_divergence_truncated_normal_distribution.png

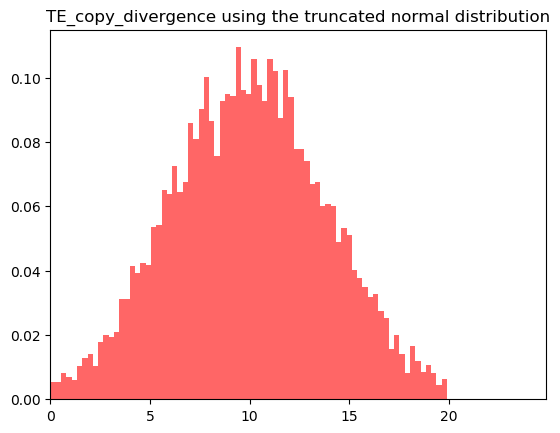

### Supplementary figures

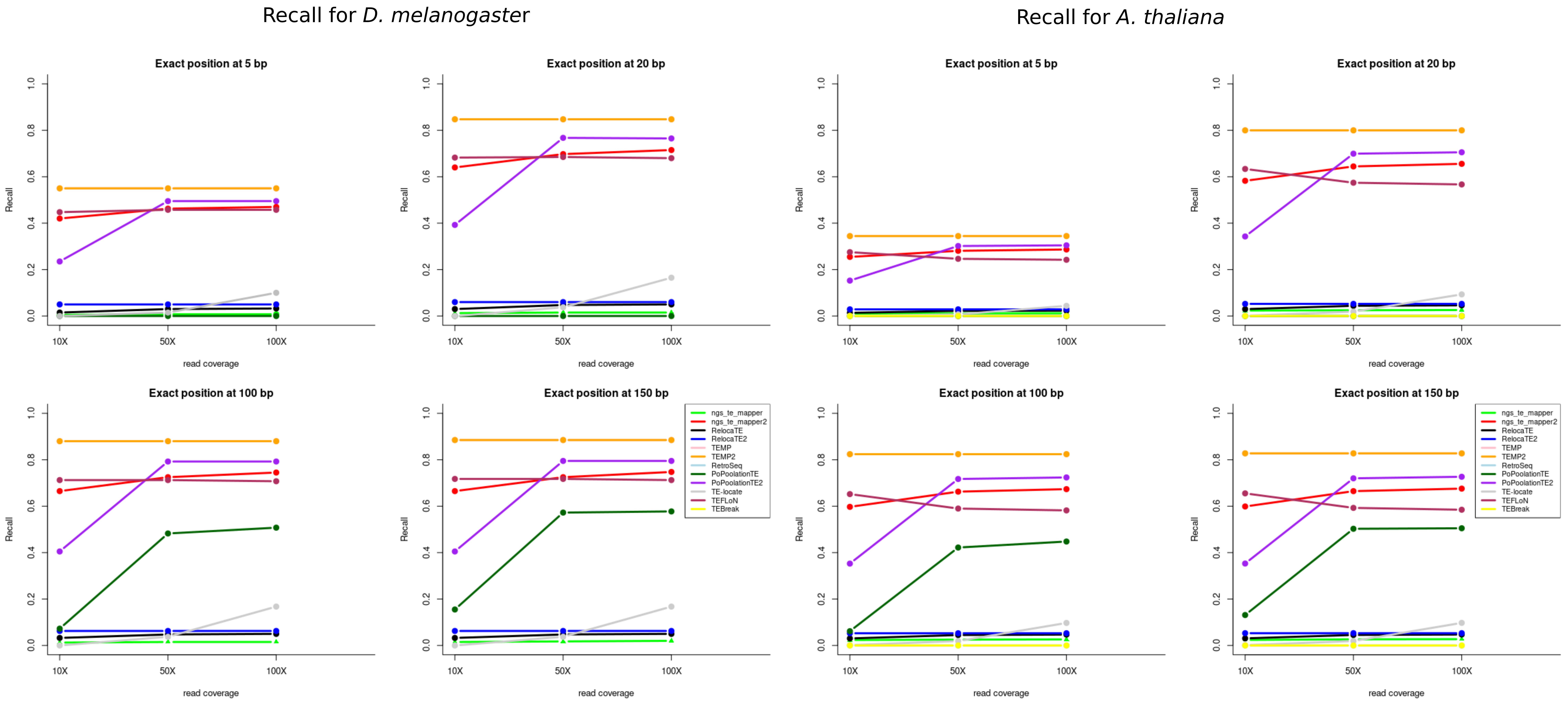

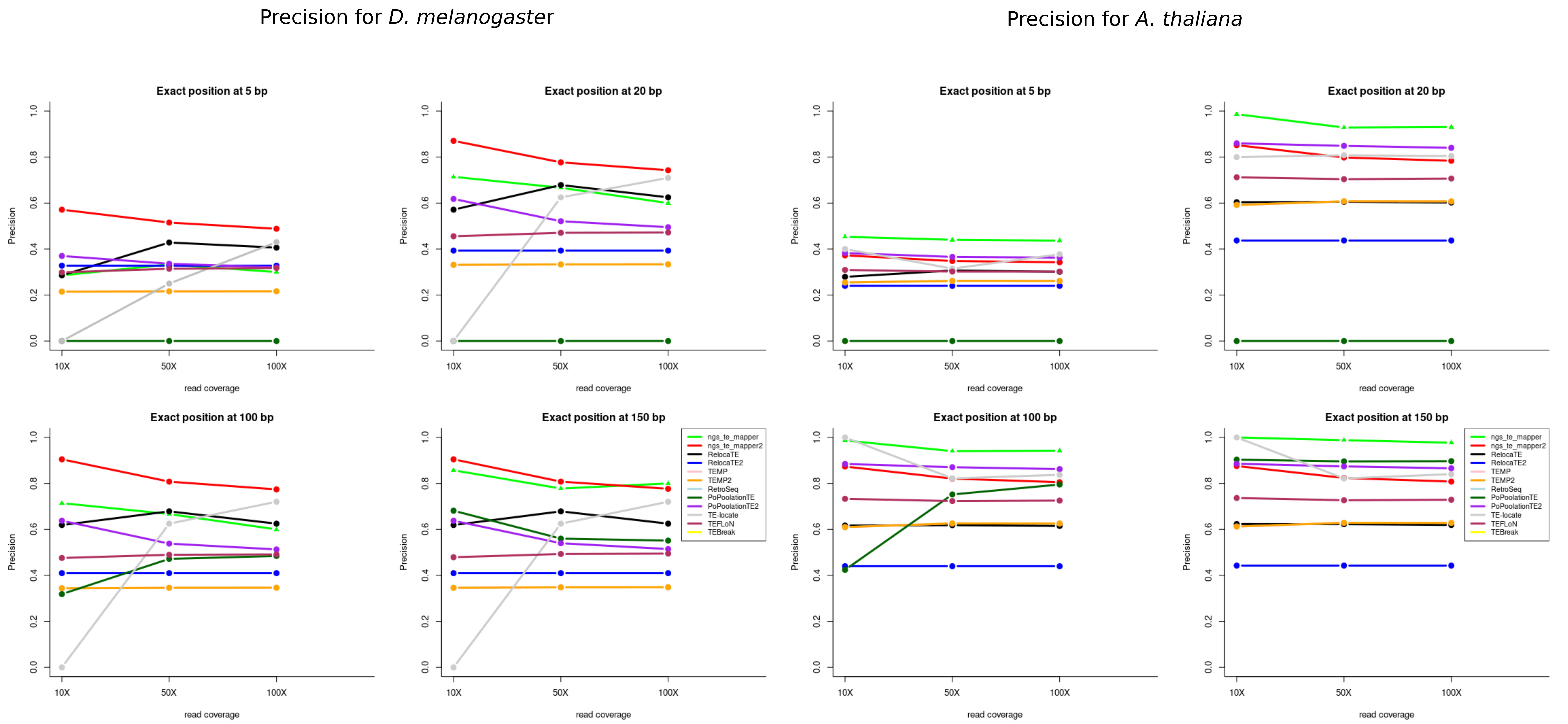

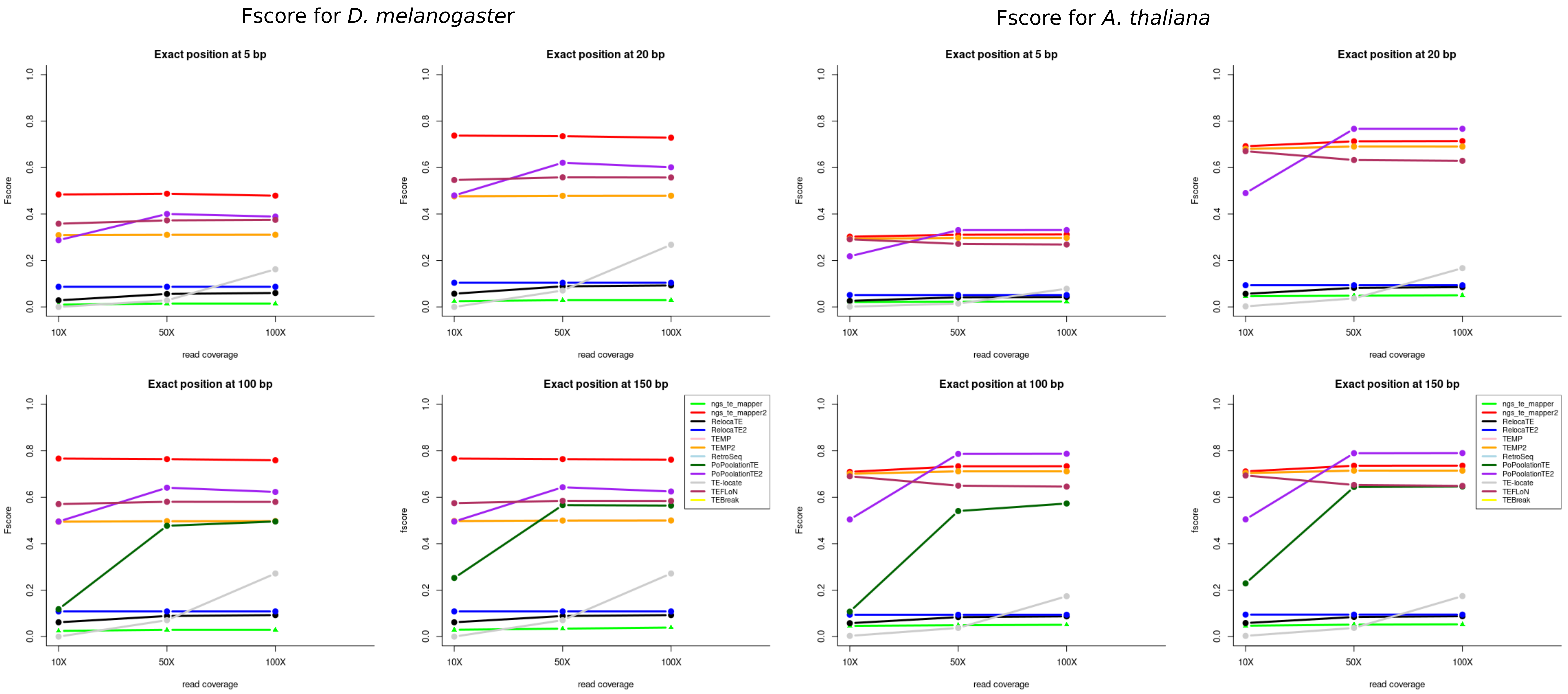

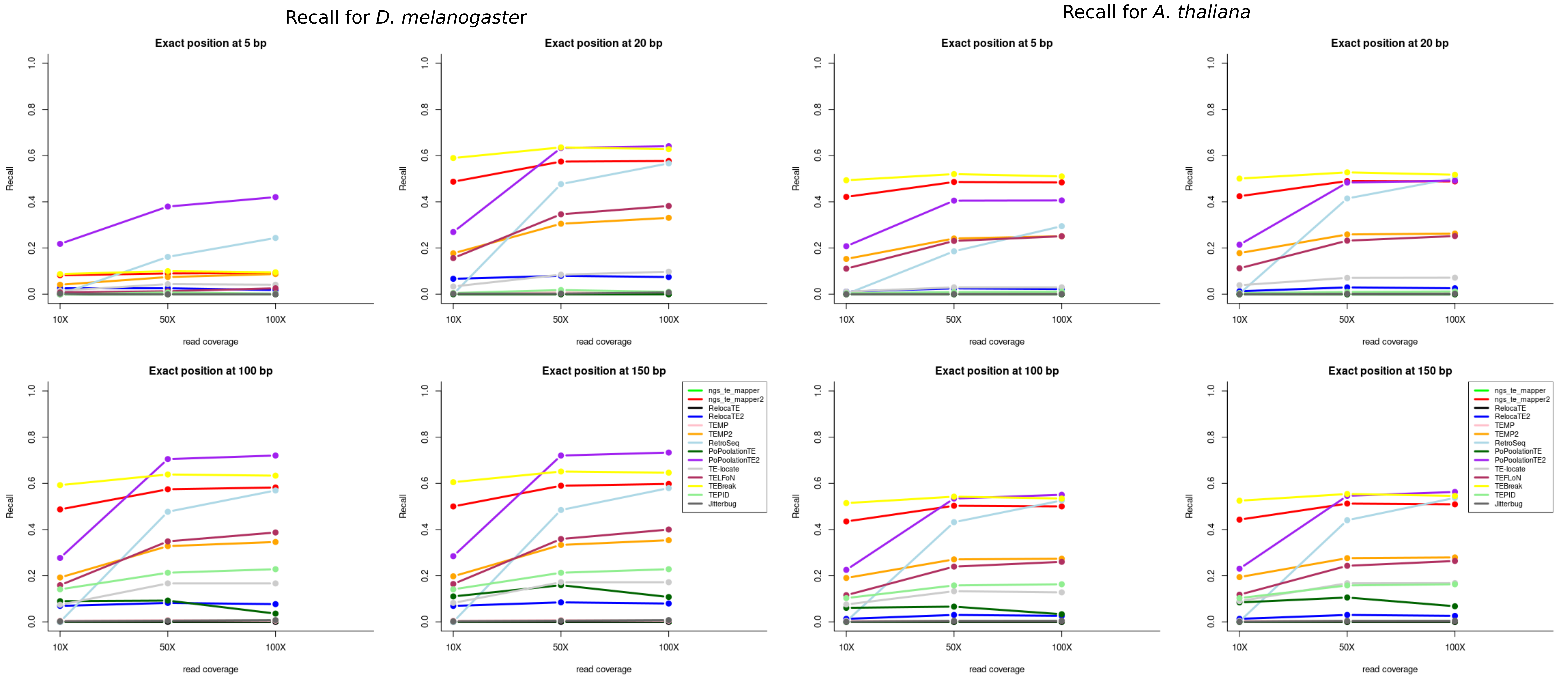

Precision for *D. melanogaster*

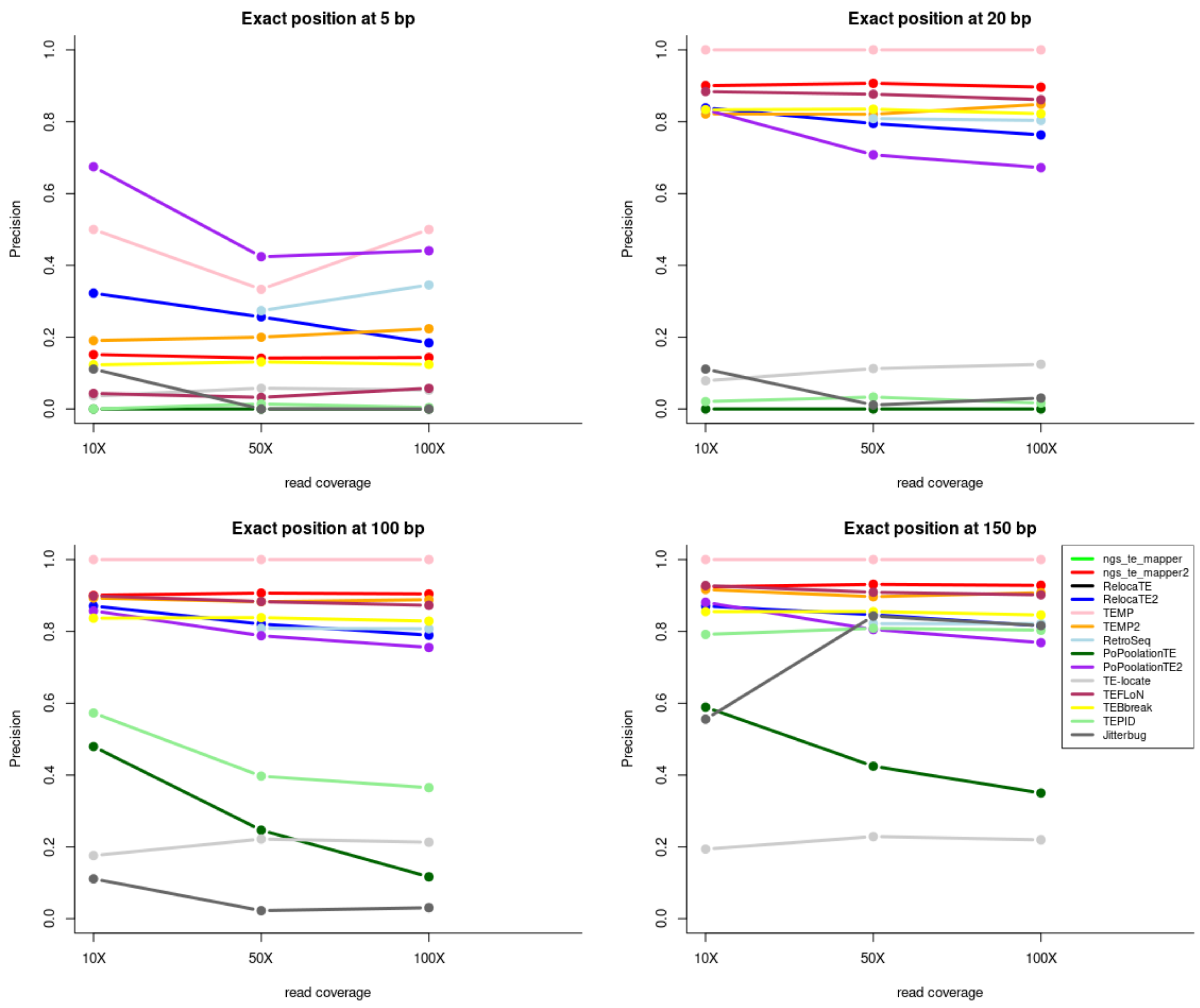


Precision for *A. thaliana*

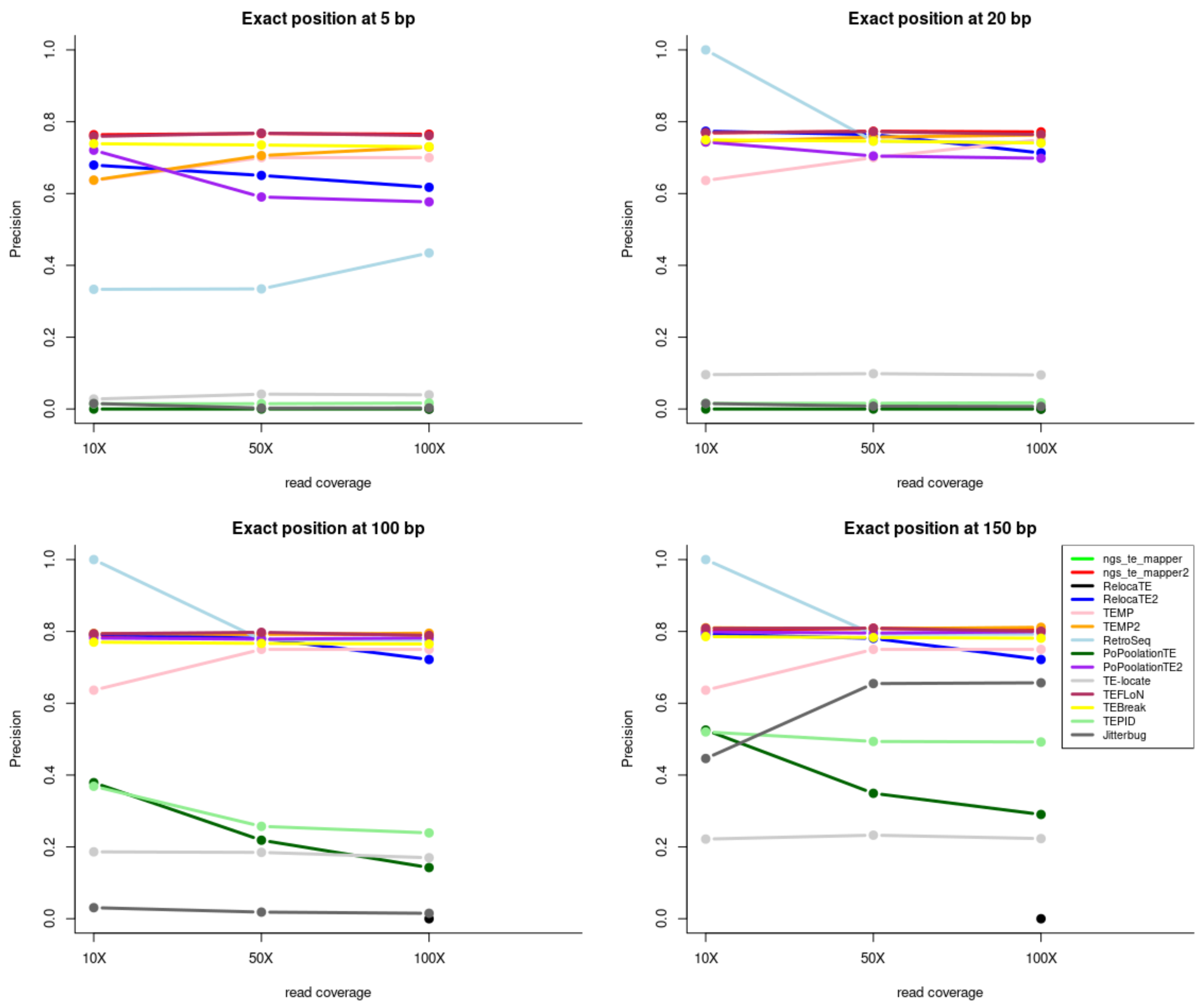

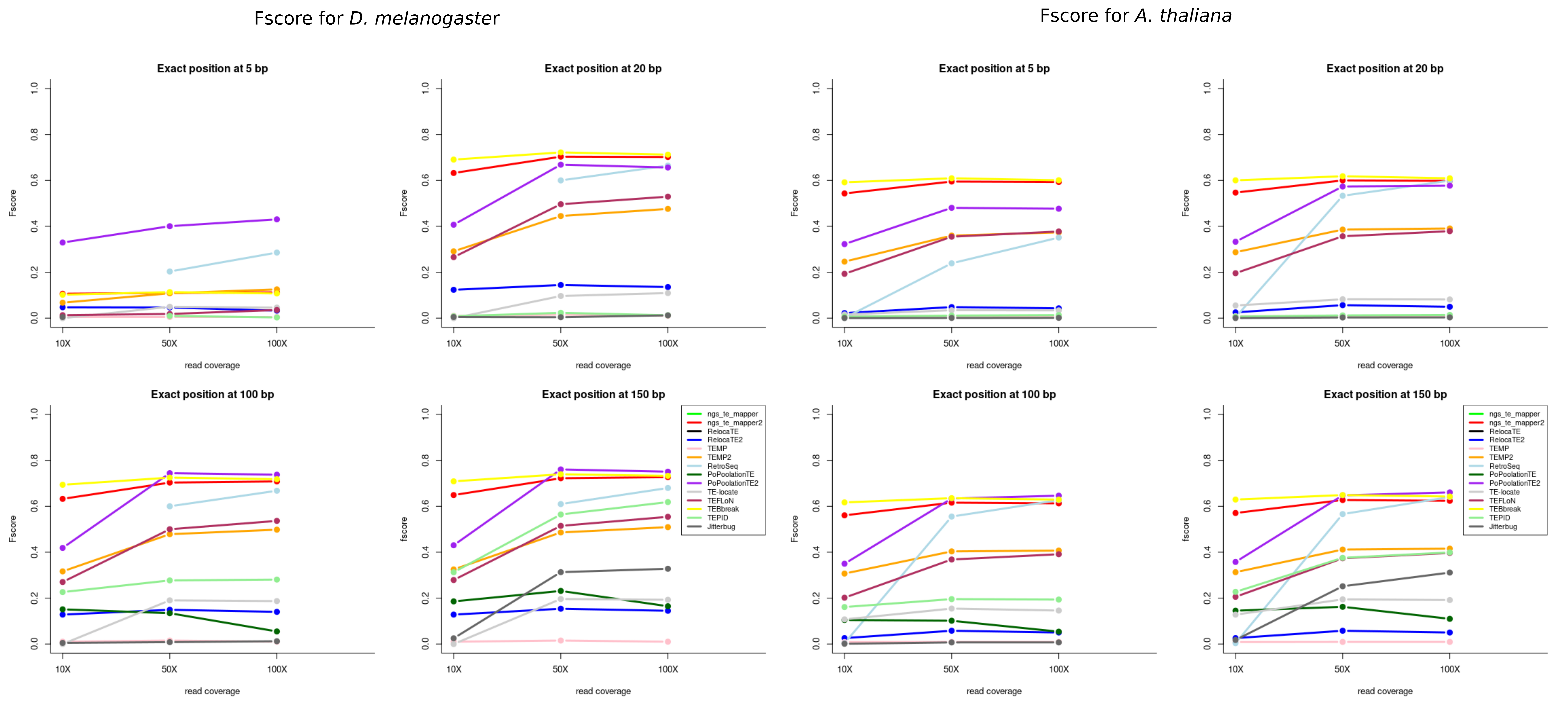

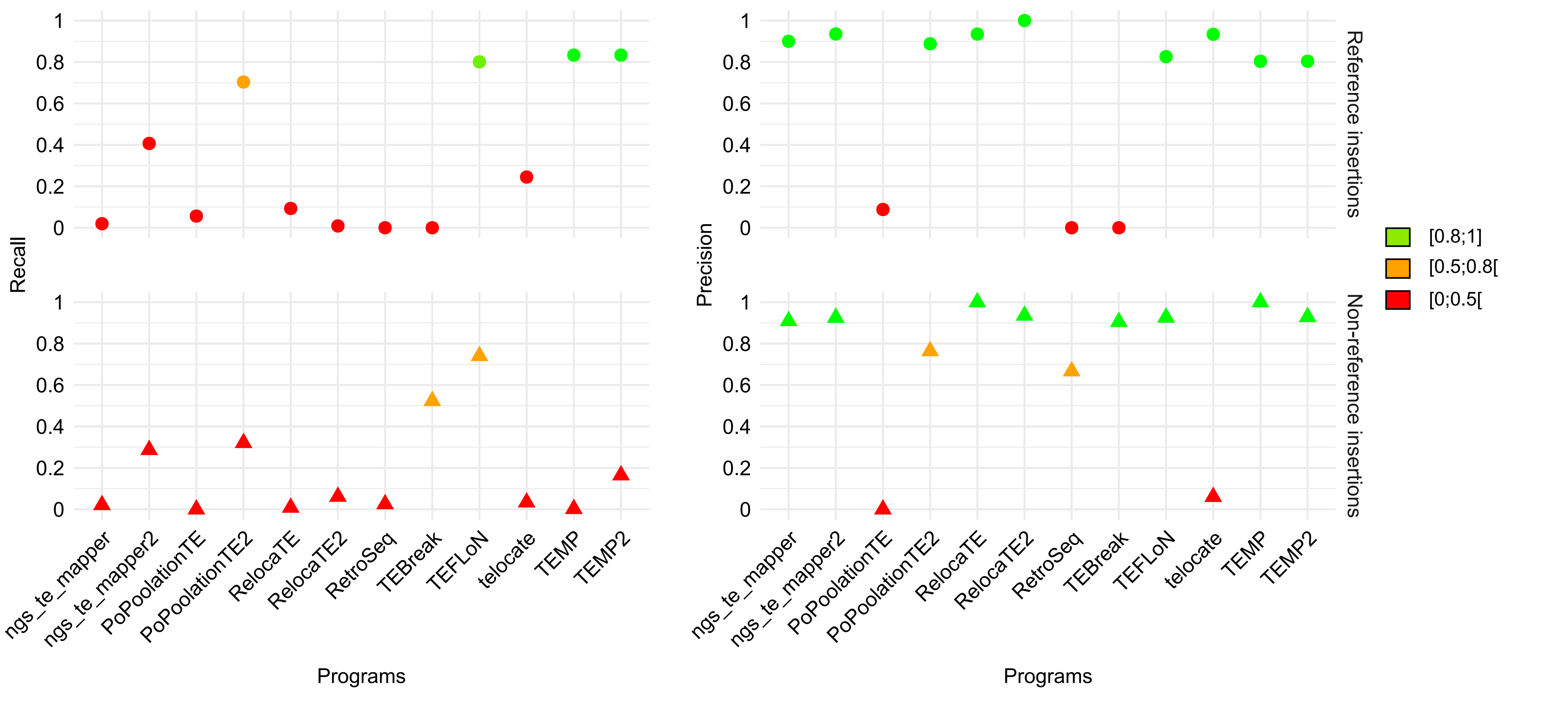

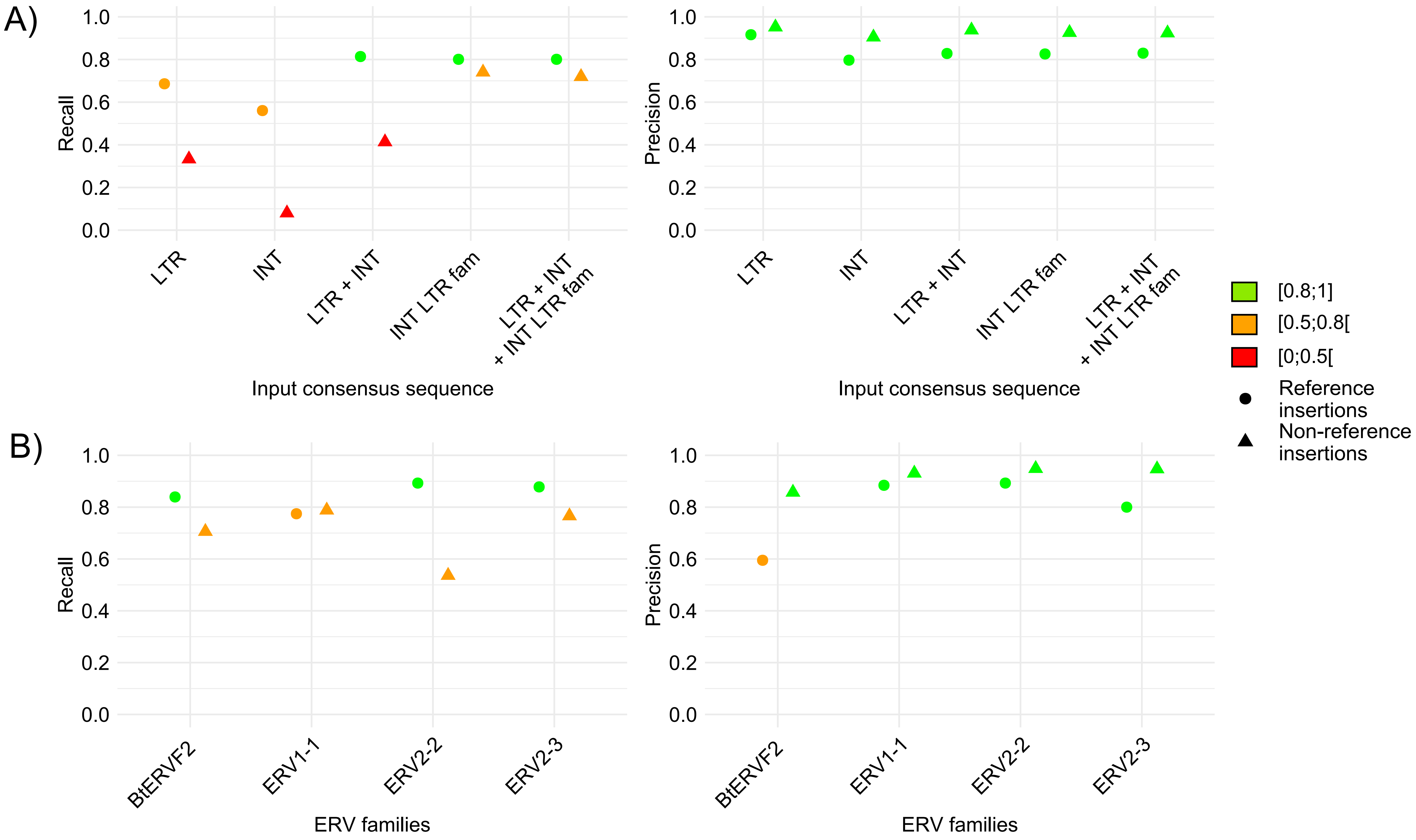

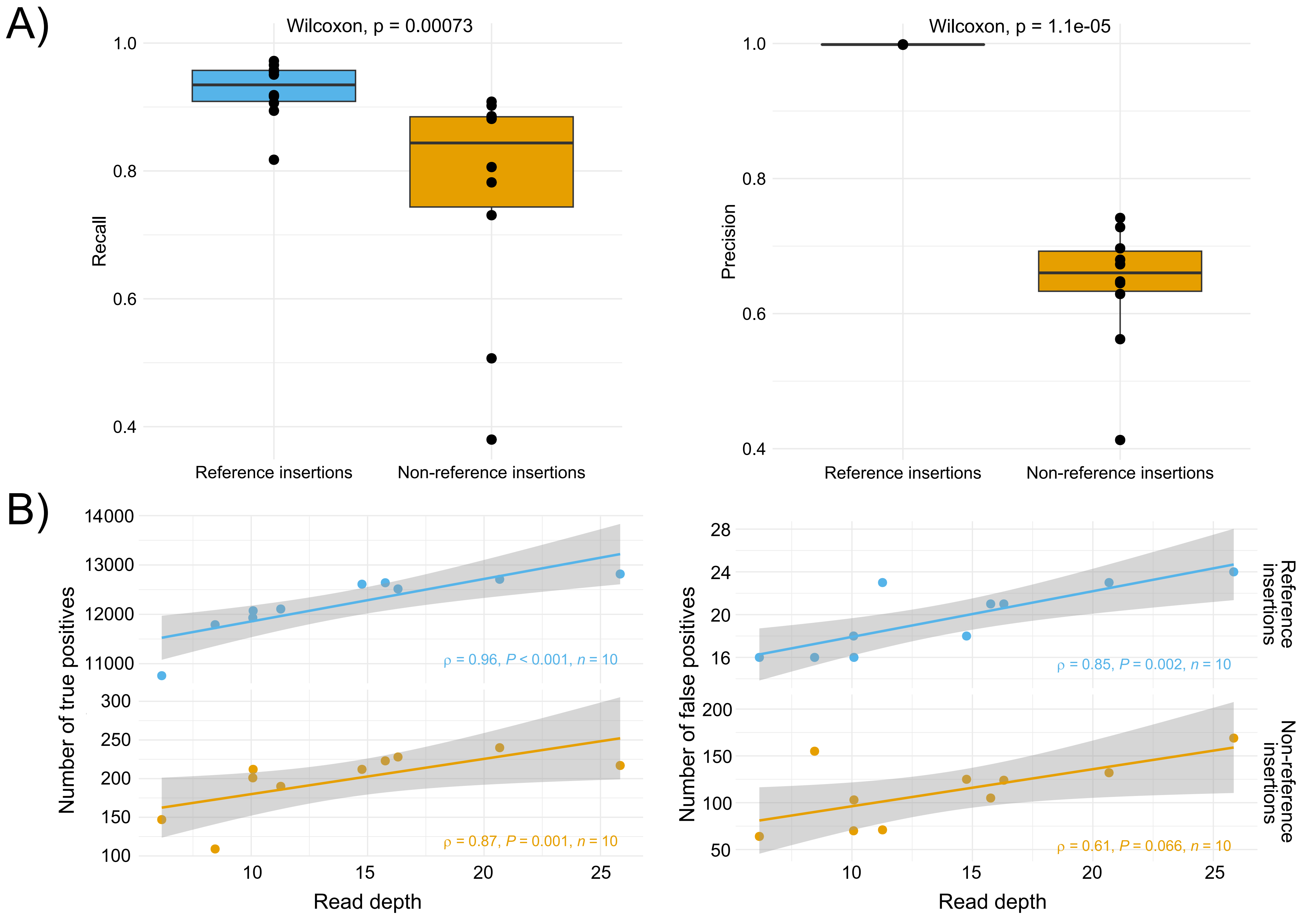
