## Supplementary file 2 for "Particular sequence characteristics induce bias in the detection of polymorphic transposable element insertions"

## Obtain long read VCF file

The VCF file for each sample was filtered to extract only insertions corresponding to our ERV families of interest

#extract the variant sequences

bcftools query -f '%CHROM\t%POS\t%ID\t%REF\t%ALT\n' ${sample}_variants.vcf
| awk -F'\t' '!/^#/' | {
    while IFS=$'\t' read -r chrom pos id ref alt; do
        if [[ $id == *INS* ]]; then
            printf ">%s\n%s\n" "$id" "$alt" >>  ${sample}.fasta
        elif [[ $id == *DEL* ]]; then
            printf ">%s\n%s\n" "$id" "$ref" >>  ${sample}.fasta
        fi
    done
} 

#perform a blast to identify only the ERV insertions

blastn -query ${sample}.fasta -db consensus_BT_repbase.fasta -out blast_ERV_${sample}.txt -outfmt 6 -evalue 1e-15 

#keep only best hits with more than 80% of identity and a size > 200bp

awk '!seen[$1]++' blast_ERV_${sample}.txt ­| awk '{OFS="\t"; if ($3>80 && $4 > 200) {print $0}}' | cut -f1 > blast_besthit_ERV_${sample}.txt
awk ‘BEGIN {while (getline id < " blast_besthit_ERV_${sample}.txt ") {keep[id] = 1}}{if ($0 ~ /^#/) {print $0} else if ($3 in keep) {print $0}}' ${sample}_variants.vcf > ERV_${sample}_variants.vcf

#extract only insertions from the vcf

awk -F'\t' '!/^#/ {split($8, info, ";") if (info[1] == "SVTYPE=INS") {print $0}}' ERV_${sample}_variants.vcf > INS_ERV_${sample}_variants.vcf

## Obtain Short read TEFLoN BED files

The TEFLoN output genotypes.txt files were converted in BED files. Only insertion with start and end coordinates were conserved and the reference and non-reference insertions were separated into different files. 

awk -F'\t' 'BEGIN { OFS = "\t" }{if ($2 > $3) {temp = $2; $2 = $3; $3 = temp} print $1, $2, $3, $7, $4, $6}' ${sample}.genotypes.txt > ${sample}.genotypes.bed
awk -F'\t' 'BEGIN{OFS="\t"} $7 == "-" && $2 != "-" && $3 != "-" { print $0 }'  ${sample}.genotypes.bed  > non-ref_${sample}.genotypes.bed
awk -F'\t' 'BEGIN{OFS="\t"} $7 != "-" && $2 != "-" && $3 != "-" { print $0 }'  ${sample}.genotypes.bed  > ref_${sample}.genotypes.bed
